# Neurexins Regulate GABA Co-release by Dopamine Neurons

**DOI:** 10.1101/2021.10.17.464666

**Authors:** Charles Ducrot, Gregory de Carvalho, Benoît Delignat-Lavaud, Constantin V.L. Delmas, Nicolas Giguère, Sriparna Mukherjee, Samuel Burke-Nanni, Marie-Josée Bourque, Martin Parent, Lulu Y. Chen, Louis-Éric Trudeau

## Abstract

Midbrain dopamine (DA) neurons are key regulators of basal ganglia functions. The axonal domain of these neurons is highly complex, with a large subset of non-synaptic release sites and a smaller subset of synaptic terminals from which glutamate or GABA are released. The molecular mechanisms regulating the connectivity of DA neurons and their neurochemical identity are unknown. Here we tested the hypothesis that the trans-synaptic cell adhesion molecules neurexins (Nrxns) regulate DA neuron neurotransmission. Conditional deletion of all Nrxns in DA neurons (DAT::Nrxns KO) showed that loss of Nrxns does not impair the basic development and ultrastructural characteristics of DA neuron terminals. However, loss of Nrxns caused an impairment of DA transmission revealed as a reduced rate of DA reuptake following activity-dependent DA release, decreased DA transporter levels, increased vesicular monoamine transporter expression, and impaired amphetamine-induced locomotor activity. Strikingly, electrophysiological recording revealed an increase of GABA co-release from DA neuron axons in the striatum of the KO mice. These findings suggest that Nrxns act as key regulators of DA neuron connectivity and DA-mediated functions.

**Highlights:**

- The study provides the first direct evidence of the role of neurexins in dopaminergic neurons.
- The synaptic adhesion molecules, neurexins, are not required for maintaining the structure of dopamine neuron terminals.
- Neurexins regulate dopaminergic neurotransmission through regulation of dopamine reuptake, impacting amphetamine-induced locomotion.
- Deletion of Nrxns in DA neurons causes a region-specific increase of GABA release by DA neurons.

## Introduction

Dopamine (DA) neurons from the ventral tegmental area (VTA) and substantia nigra pars compacta (SNc) project densely to the ventral striatum (VS) and to the dorsal striatum (DS), respectively (Descarries et al., 1980; Matsuda et al., 2009) and are critical regulators of basal ganglia functions, motivation, and cognition (Schultz, 2007; Surmeier et al., 2014). The connectivity of the DA system is predominantly non-synaptic (Descarries et al., 2008; Ducrot et al., 2021), with a majority of DA-releasing terminals not located in close apposition to a postsynaptic domain (Caille et al., 1996; Descarries et al., 1996; Descarries and Mechawar, 2000; Descarries et al., 2008; Ducrot et al., 2021). A smaller synaptic subset of DA neuron terminals has the ability to co-release glutamate or GABA (Sulzer et al., 1998; Dal Bo et al., 2004; Mendez et al., 2008; Stuber et al., 2010; Tritsch et al., 2012; Tritsch et al., 2016). The molecular mechanisms underlying the formation and regulation of non-synaptic terminals is presently undetermined. Interestingly, a growing body of work has highlighted the functional role of trans-synaptic cell adhesion molecules such as the neurexins (Nrxns) and neuroligins (NLs) in orchestrating synaptic functions and plasticity at synaptic terminals (Zhang et al 2015, Chen et al, 2017).

Nrxns are presynaptic cell adhesion molecules that were identified as α-latrotoxin receptors (Ushkaryov et al., 1992). In mammals, Nrxns are expressed in two principal forms: longer α-neurexins isoforms and shorter ß-neurexins isoforms (Tabuchi and Sudhof, 2002). The Nrxn proteins on axon terminals interact with postsynaptic NL proteins and have been shown to regulate synapse formation and function (Ichtchenko et al., 1995; Graf et al., 2004; Ko et al., 2009). NLs only bind to Nrxns, whereas Nrxns have large numbers of splice variants with differential binding affinities with multiple postsynaptic partners. Several key studies using a strategy of conditional Nrxns deletion in mice demonstrated that Nrxns regulate neurotransmission through different mechanisms in a cell-type specific manner (Chen et al., 2017; Luo et al., 2020; Luo et al., 2021).

Despite the functional importance of DA in the brain, a limited number of studies have until now explored the molecular mechanisms underlying the unique nature of the axonal arbor and connectivity of DA neurons (Liu et al., 2018; Robinson et al., 2019; Banerjee et al., 2020; Banerjee et al., 2021; Delignat-Lavaud et al., 2021; Ducrot et al., 2021) and none examined the potential role of Nrxns in these neurons. Here we tested the hypothesis that Nrxns play a key role in regulating the connectivity and functions of DA neurons by deleting all Nrxns in these cells. We crossed DAT-IRES-Cre mice with Nrxn123α/ß floxed mice [Nrxn123 Triple conditional KO mice (Chen et al., 2017)]. We found that the density and the ultrastructure of synapses established by DA neurons were not affected. However, electrophysiological recordings showed an increase in GABA release from DA terminals in the ventral but nor dorsal striatum in KO mice, suggesting a region-specific negative regulatory role of Nrxns on GABA co-transmission. DA signaling was also altered after loss of Nrxns, as revealed by slower DA reuptake, decreased density of DA transporter (DAT), and increased density of vesicular monoamine transporter (VMAT2), accompanied by impaired amphetamine-induced locomotion.

## Results

### Dopamine neuron survival and axonal connectivity are impaired by deletion of all Nrxns

Because Nrxns have been previously reported to influence the axonal development of neurons (Wang et al., 2019), we first examined the development of postnatal DA neurons *in vitro* as well as their basic connectivity. We deleted Nrxn 1, 2 and 3 from DA neurons by crossing Nrxn123^flox/flox^ mice with DAT-IRES-Cre mice (DAT::NrxnsKO) and prepared primary co-cultures of SNc or VTA DA neurons together with ventral striatal neurons obtained from postnatal day 0-3 pups (P0-P3), as previously described (Ducrot et al., 2021). DA neurons were identified by TH immunocytochemistry and automated epifluorescence imaging (**Fig. 1A**). Intriguingly, the global survival of DA neurons in DAT::NrxnsKO cultures over 14 days was significantly reduced compared to WT controls (**Fig. 1B**) (two-way ANOVA, main effect of genotype; F(1, 32)=5.00, *p*=0.032). However, we did not detect any genotype or region-specific changes in neurite development, quantified as the number and length of TH-positive neurites (**Fig. 1C** and **1D**).

**Figure 1.**
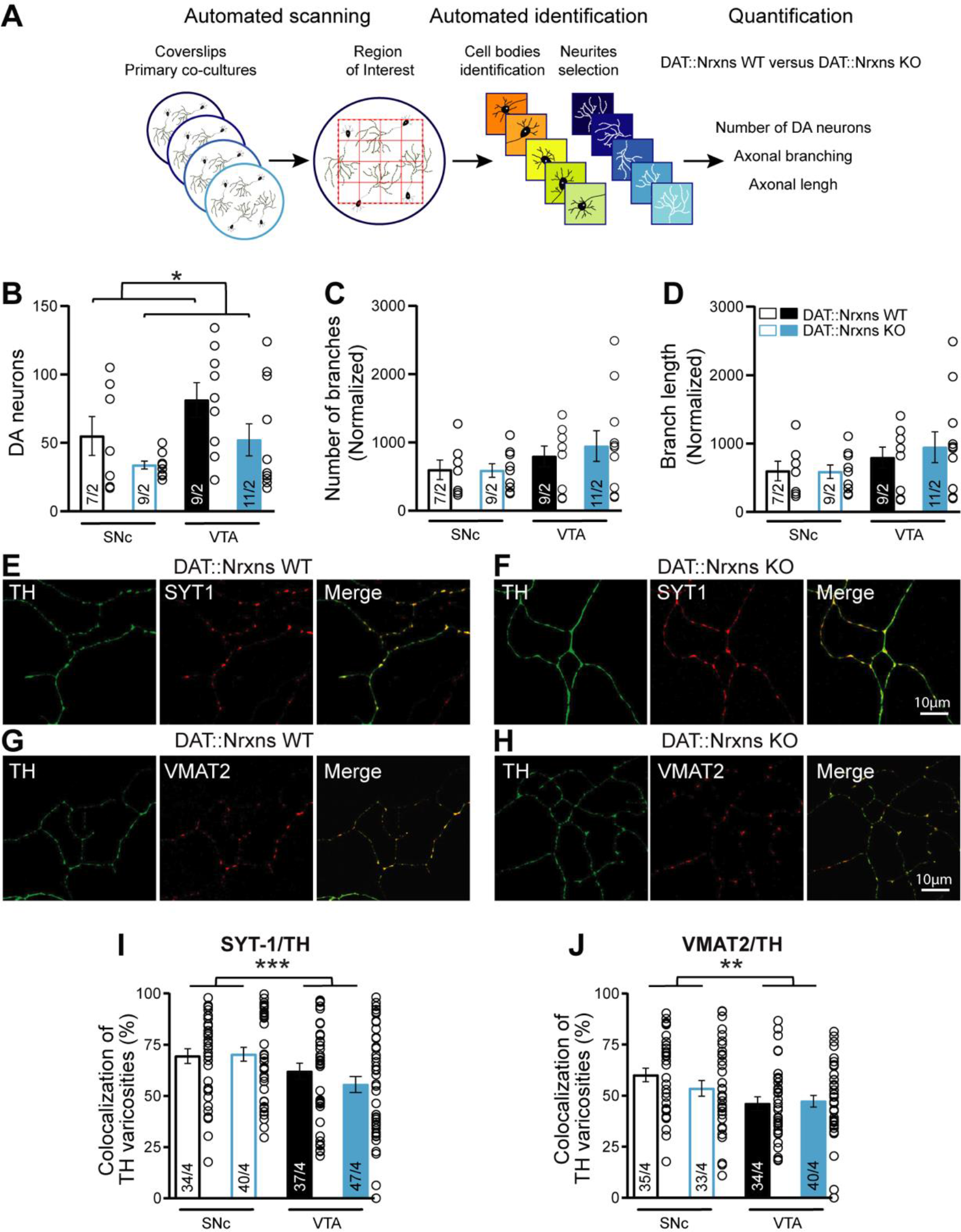
Normal axonal growth in DA neurons after conditional deletion of all neurexins. **A** - Illustration of the method used to evaluate and quantify the survival and growth of DA neurons lacking all neurexins. **B** - Quantification of DA neuron survival, assessed by their number per coverslip at 14DIV (SNc WT=55.00 ± 14.18; SNc KO=33.78 ± 2.84; VTA WT=81.33 ± 12.62; VTA KO=52.18 ± 11.72). **C** and **D** - Evaluation of neurite development in cultured DA neurons, assessed by quantifying the number of TH-positive processes (**C**) and their length (**D**) (*Branch length*: SNc WT=19086 ± 1749μm versus SNc KO=14950 ± 2068 μm, VTA WT=34721 ± 9282 μm versus VTA KO=30593 ± 4715 μm*; Branch number*: SNc WT= 598.9 ± 143.3 versus SNc KO=588.2 ± 99.37, VTA WT=795.4 ± 152.6 versus VTA KO=945.7 ± 225.7). **E and F**- Photomicrographs illustrating the distribution of TH/Syt-1 positive axonal varicosities along the axonal domain of a SNc DA neuron from DAT::NrxnsWT and DAT::NrxnsKO mice (SNc WT=60.10 ± 3.29 versus SNc KO=53.58 ± 3.81; VTA WT=46.11 ± 3.34 versus VTA KO=47.30 ± 2.86. **G and H-** Photomicrographs illustrating the distribution of TH/VMAT2 positive axonal varicosities along the axonal domain of a SNc DA neuron from DAT::NrxnsWT and DAT::NrxnsKO mice. **I-** Summary graph showing the proportion of TH-positive terminals containing Syt1. **J**- Summary graph showing the proportion of TH-positive terminals containing VMAT2. For axonal growth assessment: n=7-11 coverslips from 2 different neuronal co-cultures. For Syt1/VMAT2 quantifications: n=33-44 axonal fields from 4 different neuronal co-cultures. The number of observations represent the number of fields from individual neurons examined. For all analyses, the plots represent the mean ± SEM. Statistical analysis was carried out by two-way ANOVAs followed by Šidák’s corrections (*p < 0.05; **p < 0.01; ***p < 0.001; ****p < 0.0001).

In a second set of analyses, we also examined some of the characteristics of axon terminals established by DAT::NrxnsKO and DAT::NrxnsWT DA neurons (**Fig. 1E to 1H**). After double-labelling for TH and for Syt1, the ubiquitous exocytosis Ca^2+^ sensor, fields containing axonal arbors were examined for colocalization analysis. Our results show that the proportion of dopaminergic varicosities containing Syt1 was significantly lower in VTA DA neurons compared to SNc DA neurons as previously observed (Ducrot et al., 2021) (**Fig. 1E, 1F, 1I**; two-way ANOVA, main effect of region; F(1, 143)=13.83, *p*=0.0003), with no effect of genotype. Similarly, double labelling for VMAT2, the transporter responsible for vesicular packaging of DA, showed that the proportion of dopaminergic varicosities containing VMAT2 was significantly lower for VTA neurons compared to SNc neurons (**Fig. 1G, 1H, 1J**; two-way ANOVA, main effect of region (F(1, 138)=9.33, *p*=0.0027), with no effect of genotype. We conclude that, although loss of Nrxns may decrease the resilience of DA neurons, it does not alter the intrinsic capacity of these neurons to develop an axonal domain and to establish neurotransmitter release sites.

A subset of terminals along the complex axonal arbor of DA neurons has the capacity to release glutamate or GABA (Sulzer et al., 1998; Dal Bo et al., 2004; Mendez et al., 2008; Stuber et al., 2010; Tritsch et al., 2012; Tritsch et al., 2016). We hypothesized that deletion of Nrxns could alter the formation of excitatory or inhibitory synapses by SNc and VTA DA neurons. To test this, we co-cultured DA neurons with striatal neurons, and examined dopaminergic axon terminals in close proximity to postsynaptic organizers associated with glutamate (PSD95) and GABA (gephyrin) synapses (Craig et al., 1996; Kornau et al., 1997). We observed that loss of Nrxns reduced the proportion of SNc, but not VTA, DA neuron terminals colocalized with PSD95 (**Fig. 2A, 2B, 2C**; two-way ANOVA, main effect of region, F(1, 139)=7.65, *p*=0.0065; Sidak’s multiple comparison test, SNc WT vs SNc KO DA neurons, *p*=0.035). We also detected a significant decrease of the proportion of SNc DA neuron terminals colocalizing with gephyrin (**Fig. 2D**; two-way ANOVA, F(1, 115)=4.53; *p*=0.035; Sidak’s multiple comparison test, SNc WT vs SNc KO DA neurons, *p*=0.04). No change was observed for VTA DA neurons. In previous work, we showed that most bassoon-positive DA neuron terminals are near target cells and most likely to be found at synapses (Ducrot et al., 2021). Thus, we evaluated the proportion of DA neuron terminals expressing the glutamatergic active zone protein bassoon. We found a significant decrease in the proportion of SNc DA terminals that were bassoon positive (**Fig. 2E**; two-way ANOVA, main effect of region, F(1, 163)=5.42, *p*=0.021, and globally, there was a reduced proportion of terminals that were bassoon positive in KO DA neurons (main effect of genotype, F(1, 163)=4.32, *p*=0.039). This effect was selective for SNc DA neurons as no difference was found for VTA DA neurons (**Fig. 2E**; Sidak’s multiple comparison test, SNc WT vs SNc KO DA neurons, *p*=0.049). We conclude that these transsynaptic proteins regulate synaptic organization in a region-specific manner, compatible with previous results (Luo et al., 2020).

**Figure 2.**
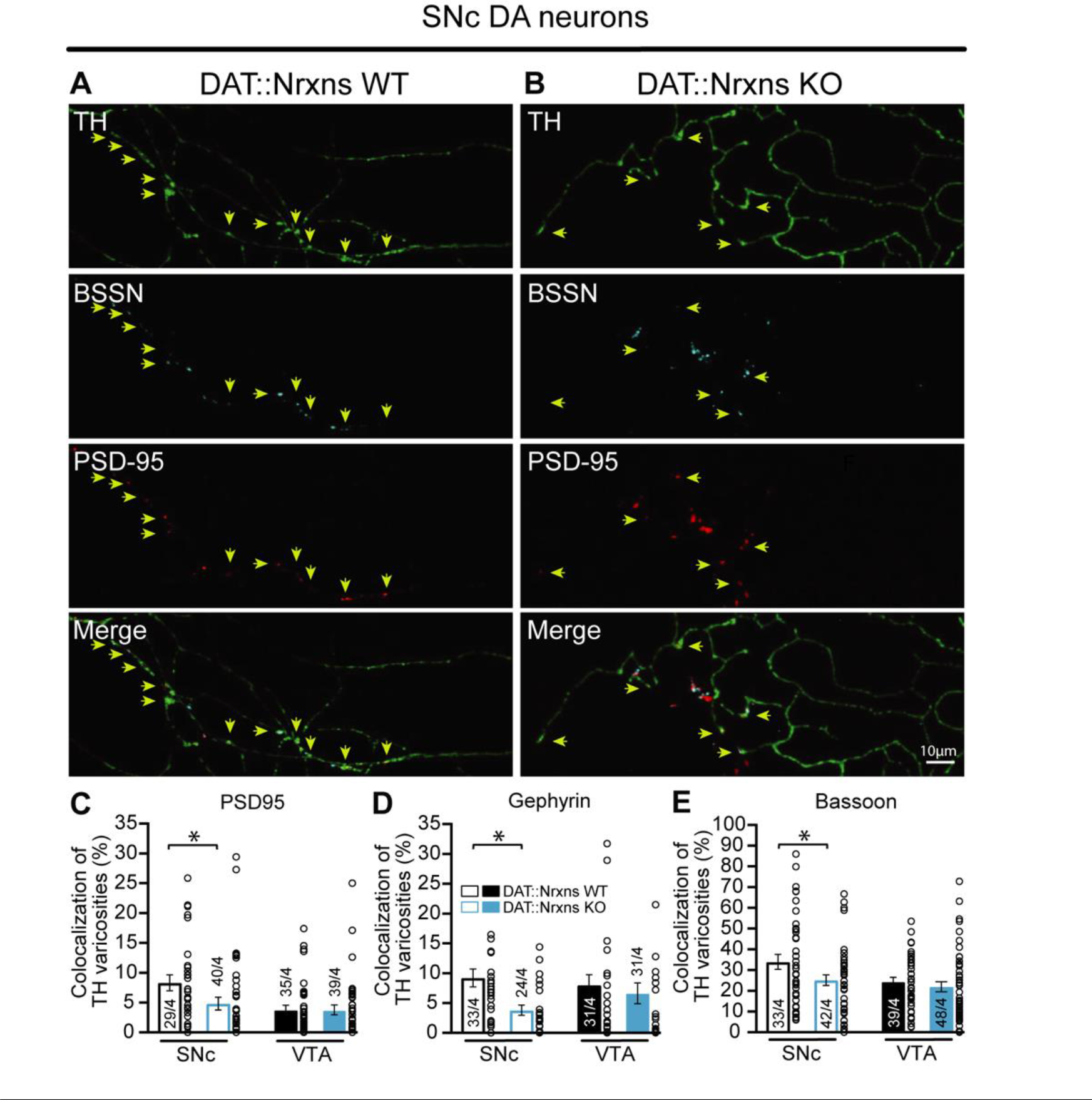
The proportion of synapses established by cultured SNc DA neurons is reduced after conditional deletion of all neurexins. **A and B-** Representative images illustrating TH-positive varicosities in the axonal arbor of a SNc DA neuron from DAT::NrxnsWT and DAT::NrxnsKO mice. Bassoon was expressed sparsely and colocalized with the postsynaptic marker PSD95. **C-** Bar graph representing the proportion (%) of axonal varicosities established by VTA and SNc DA neurons that are positive for TH and colocalizing with PSD95 at 14 DIV (SNc WT 8.32 ± 1.32% versus SNc KO 4.82 ± 1.08% and VTA WT 3.78 ± 0.78%; VTA KO 3.78 ± 0.81%). **D-** Bar graph representing the proportion (%) of axonal varicosities established by VTA and SNc DA neurons that are positive for TH and colocalizing with gephyrin at 14 DIV (SNc WT 9.22 ± 1.50% versus SNc KO 3.80 ± 0.83% and VTA WT 7.99 ± 1.76% versus VTA KO 6.64 ± 1.75%). **E -** Bar graph representing the proportion (%) of axonal varicosities that are positive for bassoon in VTA and SNc DA neurons from DAT::NrxnsWT and DAT::NrxnsKO mice (SNc WT 33.98 ± 3.62% versus SNc KO 25.09 ± 2.57% and VTA WT 24.42 ± 2.13% versus VTA KO 21.97 ± 2.43%). Data represent mean ± SEM. Statistical analyses were performed by two-tailed Student’s T-tests (*p<0.05; ** p<0.01; *** p<0.001).

### Nrxn123 ablation does not impair synapse ultrastructure in dopamine neurons

We next examined synaptic ultrastructure in the intact brain by transmission electron microscopy (TEM) to directly test our hypothesis that Nrxns regulate the integrity of synapses established by DA neurons. We focused on terminals in the vSTR because this area contains DA neuron release sites for both glutamate and GABA in addition to those for DA (Stuber et al., 2010; Berube-Carriere et al., 2012). Overall, we found that, irrespective of the genotype, most axonal varicosities contained synaptic vesicles and mitochondria (**Fig. 3A** and **3B**). Furthermore, TH-positive dopaminergic terminals in the vSTR of Nrxn123 KO were not different compared to Nrxn123 WT mice in terms of their overall perimeters (P) (**Fig. 3D**; unpaired t-test, t_6_=0.07; *p*=0.94), length (L) (**Fig. 3E**, unpaired t-test, t_6_=0.72; *p*=0.94), width (w) (**Fig. 3F**, unpaired t-test, t_6_=0.84; *p*=0.43), or surface area (**Fig. 3G**, unpaired t-test, t_6_=0.09, *p*=0.92).

**Figure 3.**
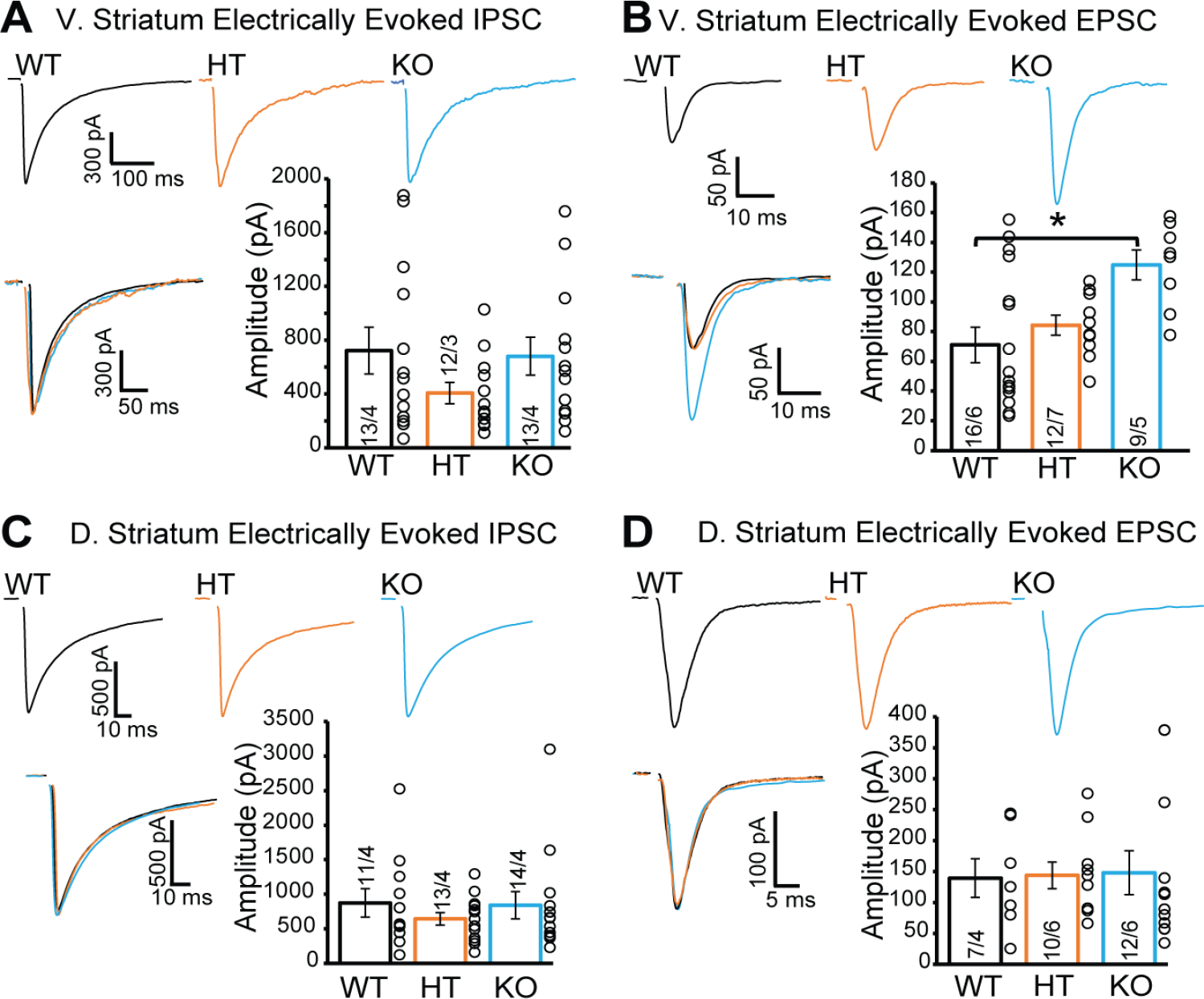
Synaptic and non-synaptic ultrastructure of DA terminals is unchanged after Nrxn123 triple knockout in dopamine neurons. **A-B** Electron micrographs showing DA neuron terminals without any PSD domain (top images) or in apposition to a PSD domain in ventral striatal tissue from DAT::NrxnsWT and KO mice. The lower micrograph represents a magnified view of the regions identified by the doted lines in the middle images. The asterisk identifies a synapse and the black arrowheads delimitate the postsynaptic domain. **C**- Schematic representation of a dopaminergic varicosity. **D**- Bar graph representing the perimeter of the DA axonal varicosity from WT and KO mice (2227 ± 81.83 nm and 1870 ± 174.8 nm, respectively), **E** and **F**- Bar graphs representing the size of the axonal varicosities, quantified as length (**E**) (897.3 ± 38.06 nm and 902.7 ± 38.06 nm respectively) and width (**F**) (468.7 ± 38.06 nm and 375.2 ± 22.02 nm, respectively). **G**- Bar graphs showing the surface area of DA neuron varicosities from WT and KO animals (323537 ± 45861 nm2 and 317887 ± 40227 nm2, respectively). **H**- Bar graphs representing the PSD domain size (232.8 ± 23.40 nm and 197.1 ± 35.71 nm, respectively for WT and KO mice). For all analyses, WT=101 and KO=189 axonal varicosities from 4 different mice for each genotype. For all analyses, plots represent the mean ± SEM. Statistical analyses were carried out by unpaired t-tests (*p < 0.05; **p < 0.01; ***p < 0.001; ****p < 0.0001).

In addition, the propensity of these terminals to make contact with a postsynaptic density (PSD) domain was unchanged in DAT::NrxnsKO mice. The synaptic incidence of TH-positive terminals was 6.34% (12/189 examined varicosities) for DAT::NrxnsKO mice and 4.95% (5/101 examined varicosities) for control mice (not shown), a low proportion in line with previous work (Stuber et al., 2010; Berube-Carriere et al., 2012). The size of the PSD was also unchanged (**Fig. 3H**, unpaired t-test, t15=0.61, *p*=0.54). Together, these results show that loss of Nrxns123 does not impair the basic ultrastructure of DA neuron neurotransmitter release sites in the vSTR.

### Deletion of Nrxns reduces amphetamine-induced locomotion without affecting basal motor activity or coordination

Although we did not observe an overall structural change in DA neuron terminals in Nrxn123KO mice, a region-specific change in both excitatory and inhibitory synaptic connections established by DA neurons was detected. Because DA neurons are key regulators of movement, motivation, and reward-dependent learning and several studies using mouse lines with impaired DA transmission reported deficits in basal or psychostimulant-evoked locomotion and learning on the accelerating rotarod (Zhou and Palmiter, 1995; Ogura et al., 2005; Birgner et al., 2010), we therefore examined DA-dependent behaviors in Nrxn123 KO mice. In the first series of experiment, we evaluated motor coordination and learning using the accelerating rotarod task with two different protocols (**Fig. 4A**). The first protocol evaluated the rate of learning to perform this task over a total of 9 sessions during 3 days, with two sessions performed on the first day, three sessions the second day, and four sessions on the third day, with a speed of rotation accelerating from 4 to 40 rpm over 10 min. The measure of latency to fall did not reveal a significant difference between the genotypes, with all groups showing a comparable increase in performance (**Fig. 4B**, two-way repeated measures ANOVA, F(2, 22)=1.59, *p=*0.22). Similar results were obtained when evaluating the progression of the performance of the mice by comparing the first and last sessions, with all mice showing equivalent learning (**Fig. 4C**; two-way ANOVA, main effect of training session, F(1, 22) = 48.71, *p*˂0.0001; Sidak’s multiple comparisons test, S1 vs S9 : WT, *p*=0.022; HET, *p*=0.0002; KO, *p*=0.001; no genotype effect was observed). The speed of rotation at the end of each trial across all 9 trials was also unchanged (**Fig. 4D**; two-way repeated measures ANOVA, F(2, 22)=1.52, *p=*0.24). Furthermore, testing a separate cohort of mice with a more challenging version of the rotarod task (**Fig. 4A**), with speed of rotation accelerating from 4 to 40 rpm over 2 min, revealed that performance, calculated by the latency to fall, was not significantly different in DAT::NrxnsKO and DAT::NrxnsHET compared to DAT::NrxnsWT (**Fig. 4E**, two-way ANOVA, repeated measures, F(2, 29)=2.81, *p*=0.07). In this task, performance failed to improve over the trials, revealing a limited capacity to improve performance, as shown by comparing performance in the last session compared to the first (**Fig. 4F**; two-way ANOVA, F(1, 29) = 0.097, *p*=0.75). The speed of rotation at the end of each trial across all 9 trials (**Fig. 4G**) was similar in DAT::NrxnsKO mice compared to the control mice (two-way repeated measures ANOVA, F(2, 29)=2.54, *p*=0.09).

**Figure 4.**
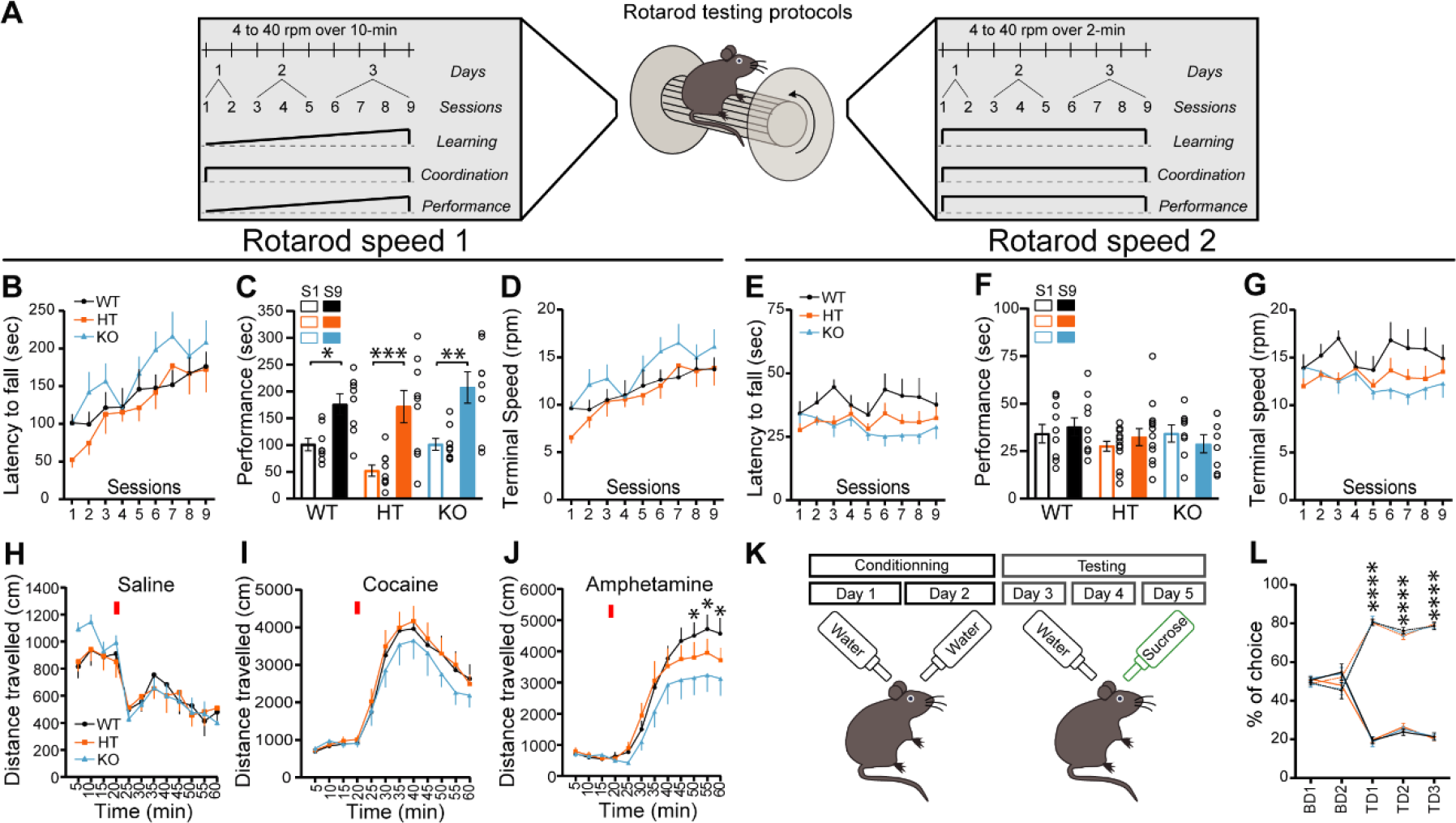
DAT::NrxnsKO mice exhibit impaired amphetamine-induced motor activity. **A-** Schematic representation of a mouse on a rotarod and the diagram of the two rotarod testing protocols. **B**- Performance on the accelerating rotarod during 9 sessions over 3 consecutive days. Latency to fall was quantified at rotation speeds from 4 to 40 rpm over 10 min. **C**- Performance of DAT::NrxnsKO, HET and WT littermate mice on the rotarod were evaluated comparing the last session and the first session for each mouse. The results show a significant improvement in performance irrespective of genotype. **D-** Quantification of the terminal speed over all the sessions shows no difference between the DAT::NrxnsKO, HET and WT littermate mice. **E**- Performance on the second accelerating rotarod task during 9 sessions over 3 consecutive days. Latency to fall was quantified at rotation speeds from 4 to 40 rpm over 2-min. **F**- Performance of DAT::NrxnsWT, HET and KO littermate mice on the rotarod was evaluated comparing the last and first sessions for each mouse. No significant improvement in performance was detected, irrespective of genotype. **G**- Quantification of the final speed over all sessions shows no difference between the DAT::NrxnsWT, HET and KO littermate mice**. H**- Basal horizontal activity in a novel environment before and after a saline injection (10ml/kg) over a total of 60-min. **I**- Horizontal activity before and after a cocaine injection (20mg/kg; 10ml/kg) over a total of 60-min. **J**- Horizontal activity before and after an amphetamine injection (5mg/kg; 10ml/kg) over 60-min, shows reduced locomotion in the DAT::NrxnsKO compared to the control mice. **K**-Schematic representation of the sucrose preference testing protocol. **L**- Quantification of sucrose preference in comparison to water consumption represented as a percentage. Initial two days: DAT::NrxnsKO CD1: 51.47 ± 1.63% vs 48.52 ± 1.63% and CD2: 54.01 ± 3.17%, vs 45.99 ± 3.17%; DAT::NrxnsHET CD1: 51.16 ± 1.81% vs 48.83 ± 1.81% and CD2: 47.90 ± 2.74%, vs 52.09 ± 2.74%; DAT::NrxnsWT CD1: 50.58 ± 1.47% vs 49.41 ± 1.47% and CD2: 54.73 ± 4.27%, vs 45.26 ± 4.27%. Results are presented as percentage of choice water/water. Following three test days: DAT::NrxnsKO TD1: 81.24 ± 2.44% vs 18.75 ± 2.44%; TD2: 74.65 ± 1.39%, vs 25.34 ± 1.39% and TD3: 78.74 ± 1.37%, vs 21.25 ± 1.37%; DAT::NrxnsHET TD1: 80.30 ± 1.39% vs 19.69 ± 1.39%; TD2: 73.57 ± 1.85%, vs 26.43 ± 1.85% and TD3: 79.68 ± 0.96% vs 20.32 ± 0.96%; DAT::NrxnsWT TD1: 80.52 ± 1.74% vs 19.47 ± 1.74%; TD2: 76.21 ± 1.75%, vs 23.78 ± 1.75% and TD3: 78.58 ± 2.00%, vs 21.41 ± 2.00%). Results are presented as percentage of choice water/water. Following three test days: DAT::NrxnsKO TD1: 81.24 ± 2.44% vs 18.75 ± 2.44%; TD2: 74.65 ± 1.39%, vs 25.34 ± 1.39% and TD3: 78.74 ± 1.37%, vs 21.25 ± 1.37%; DAT::NrxnsHET TD1: 80.30 ± 1.39% vs 19.69 ± 1.39%; TD2: 73.57 ± 1.85%, vs 26.43 ± 1.85% and TD3: 79.68 ± 0.96% vs 20.32 ± 0.96%; DAT::NrxnsWT TD1: 80.52 ± 1.74% vs 19.47 ± 1.74%; TD2: 76.21 ± 1.75%, vs 23.78 ± 1.75% and TD3: 78.58 ± 2.00%, vs 21.41 ± 2.00%). All results are presented as percentage of choice sucrose/water. For rotarod and locomotor activity experiments, 7 to 14 animals per group were used. For sucrose experiment, 7 to 86 mice per group were used). For all analyses, the plots represent the mean ±

These results suggest that deletion of Nrxn123 from DA neurons do not lead to motor coordination deficits.

General motor abilities were next evaluated using the pole test and the open field test. In the pole test (**Fig. S1A**), no difference was observed between genotypes for the time required for the mice to orient downward (**Fig. S1B)** (one-way ANOVA, F(2; 25)=0.92, *p*=0.4) and for the time required to climb down the pole (**Fig. S1C)** (one-way ANOVA, F(2; 25)=2.8, *p*=0.08). Basal locomotion in the open field over a 60-min period was also not different between genotypes (**Fig. 4H**; two-way ANOVA, F(2; 28)=0.95, *p*=0.95). We next challenged the dopaminergic system of these mice using the psychostimulants cocaine and amphetamine (Di Chiara and Imperato, 1988).

Although locomotion induced by cocaine (5mg/kg) was comparable between genotypes (**Fig. 4I**, two-way ANOVA, F(2; 26)=0.36, *p*=0.67), locomotion induced in response to amphetamine was strongly reduced in DAT::NrxnsKO mice compared to DAT::NrxnsHET and DAT::NrxnsWT mice (**Fig. 4J**, two-way ANOVA, main effect of time; F(11; 228)=45.95 and genotype F(2, 228)=9.67, *p*˂0.0001). The maximal effect was observed between 30 to 40 min post-injection (**Fig. 4J**; two-way ANOVA, Tukey’s multiple comparisons test, DAT::NrxnsKO vs WT. t=50min: *p*=0.03; t=55min: *p*=0.01; t=60min: *p*=0.02). These results suggest that loss of Nrxns in DA neurons leads to altered DA neurotransmission and DA-dependent behaviors.

Because altered DA neurotransmission is often associated with changes in states of motivation, we next examined the performance of the mice in a well-established sucrose preference task. On the initial two conditioning days (CD1 and CD2), mice of all genotypes equally licked at both bottles **(Fig. 4K, 4L)**. Similarly, during the next 3 testing days (TD1, 2 and 3), when mice were given a choice between water and sucrose, DAT::NrxnsKO, HET and WT mice showed a similar marked preference for the sucrose bottle (**Fig. 4L**; two-way ANOVA, main effect of choice F(5; 40)=498.0; Tukey’s multiple comparisons test, Test Day 1, 2, 3, water versus sucrose, all genotypes, *p*˂0.0001). These findings suggest that the response of DAT::NrxnKO mice to natural rewards was unaltered.

### Fast Scan Cyclic Voltammetry reveals normal DA release but slower reuptake and enhanced paired-pulse depression after conditional deletion of all Nrxns in DA neurons

The impaired response to amphetamine suggests a perturbation of extracellular DA dynamics or DA action on target cells. To examine this possibility, we first employed fast scan cyclic voltammetry (FSCV) to measure electrically-evoked DA overflow in acute brain slices of the ventral and dorsal striatum (vSTR and dSTR). In the dSTR, we found no difference in peak DA overflow between the DAT::NrxnsWT and DAT::NrxnsKO mice, but surprisingly, in the DAT::NrxnsHET mice, peak DA overflow was significantly lower (One-way ANOVA, F(2; 179)=7.06, *p*=0.001, Tukey’s multiple comparisons test, WT vs HET, *p*=0.002 and HET vs KO, *p*=0.006) compared to the WT and KO mice (**Fig. 5A** and **5B**).

**Figure 5.**
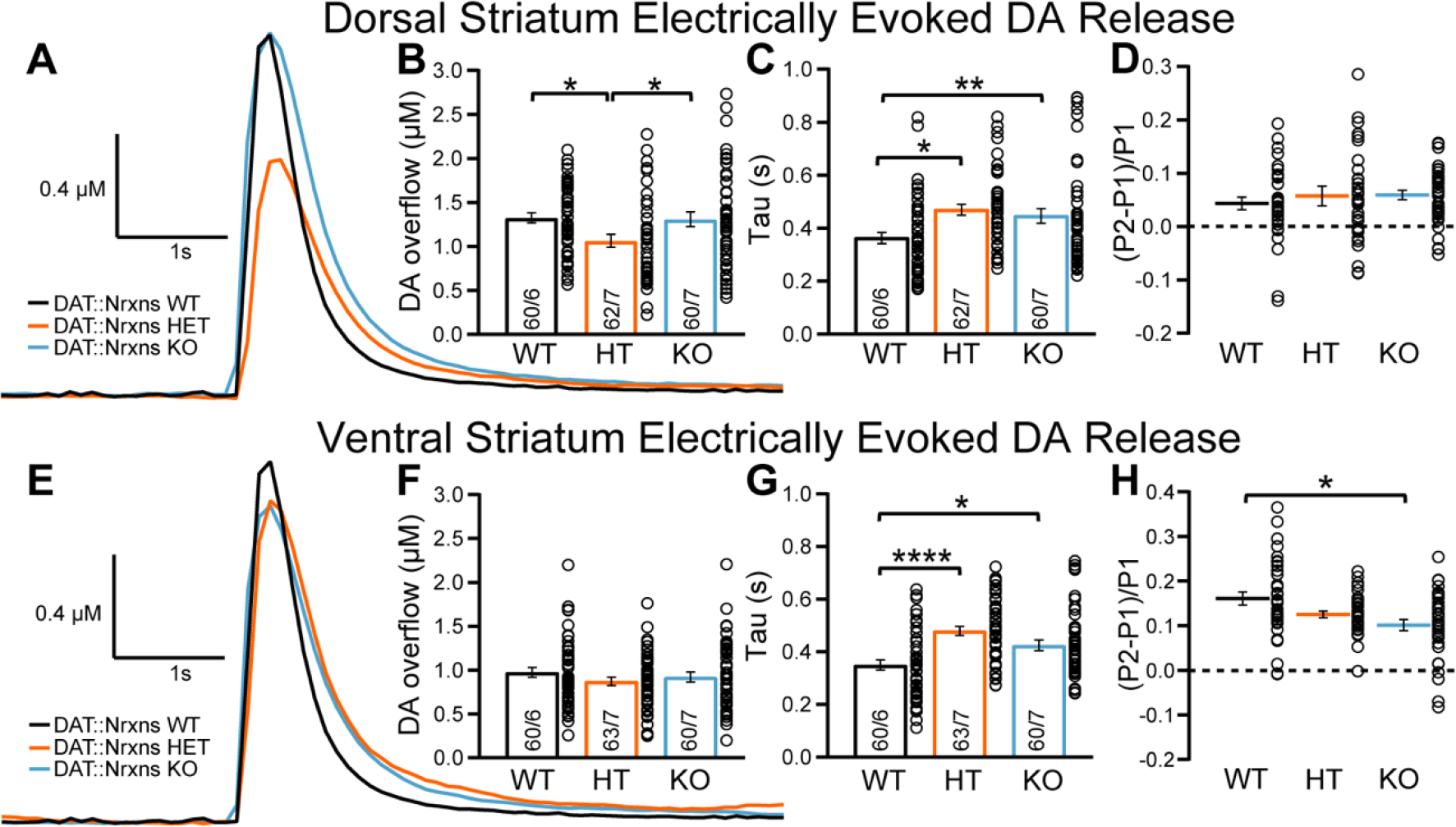
Slower dopamine reuptake in DAT::NrxnsKO mice. **A**- Representative traces of electrically evoked DA levels detected by fast-scan cyclic voltammetry in the dorsal striatum, measured in slices prepared from DAT::NrxnsWT, HET and KO mice. **B**- Bar graphs showing the average DA levels (μM) detected in the dorsal striatum (WT=1.33 ± 0.05μM; HET=1.05 ± 0.05μM and KO=1.35 ± 0.07μM). **C**- Evaluation of DA release kinetics in the dorsal striatum estimated by quantifying Tau, corresponding to reuptake efficiency (WT=0.36 ± 0.02 s; HET=0.47 ± 0.02 s and KO=0.45 ± 0.03 s). **D-** Short-term paired-pulse induced plasticity of DA overflow in dorsal striatal slices, estimated by calculating (P2-P1/P1) with an inter-pulse interval of 100 ms. The low ratio values reflect the strong paired-pulse depression seen at such release sites in acute brain slices. **E**- Representative traces of electrically evoked DA levels detected by fast-scan cyclic voltammetry in the ventral striatum. **F**- Bar graphs showing the average of DA levels (μM) detected in the ventral striatum (WT=0.98 ± 0.04μM; HET=0.88 ± 0.05μM and KO=0.98 ± 0.06μM). **G**- Evaluation of DA release kinetics in the ventral striatum, estimated by quantifying Tau (WT=0.35 ± 0.02; HET=0.47 ± 0.02 and KO=0.42 ± 0.02). **H**- Short-term paired-pulse induced plasticity of DA overflow in ventral striatal slices, estimated by calculating (P2-P1/P1) with an inter-pulse interval at 100-ms. The low ratio values reflect the strong paired-pulse depression seen at such release sites in acute brain slices. Data are presented as mean ± SEM. Statistical analyses were performed with a one-way ANOVA (*p<0.05; ** p<0.01; *** p<0.001).

An examination of the kinetics of DA overflow is often used to identify changes in DA release efficiency and reuptake (Yorgason et al., 2011). We found that DA reuptake in the dSTR was significantly slower in DAT::NrxnsKO and DAT::NrxnsHET compared to DAT::NrxnsWT mice (One-way ANOVA, main effect of genotype, F(2,140)=5.82, *p*=0.0037), as illustrated by higher Tau values (**Fig. 5A** and **5C**; Tukey’s multiple comparisons test; WT vs KO *p*=0.034; WT vs HET *p*=0.041). We also observed a similar increase in Tau for DA reuptake in the vSTR (**Fig. 5E** and **5G;** One-way ANOVA, main effect of genotype, F(2, 137)=12.17, *p*˂0.0001; Tukey’s multiple comparisons test; WT vs KO *p*=0.017; WT vs HET *p*˂0.0001).

Quantification of the rise-time of evoked DA overflow in the dSTR revealed a slower rise time in DAT::NrxnsHET mice compared to DAT::NrxnsWT mice (**Fig. S2A;** One-way ANOVA, main effect of genotype, F(2, 141)=3.23, *p*=0.42, Holm-Šidák’s multiple comparisons test, HET vs WT, *p*=0.044). No differences were found in the vSTR (**Fig. S2B)**.

Short-term plasticity of electrically evoked DA release in the striatum was examined using a paired-pulse stimulation paradigm. DA overflow in acute brain slices typically shows a large paired-pulse depression, more extensively so in the dSTR compared to the vSTR (Zhang and Sulzer, 2004; Sanchez et al., 2014; Condon et al., 2019). We found a significant difference that was selective for the vSTR. Indeed, the level of paired-pulse depression was similar in DAT::NrxnsKO mice compared to WT in the dSTR (**Fig. 5D**; One-way ANOVA; F(2, 107)=0.36, *p*=0.69). However, in the vSTR, paired-pulse depression was significantly enhanced in DAT::NrxnsKO mice compared to WT (**Fig. 5H**; One-way ANOVA; F(2, 103)=4.88, *p*=0.0094; Tukey’s multiple comparison test, *p*=0.0082). Plasticity of DA release was also examined by measuring the inhibition of DA overflow induced by the GABA_B_ agonist baclofen (10µM) (Lopes et al., 2019). A recent study demonstrated a role for Nrxns in the regulation of the expression and location of presynaptic GABA_B_ receptors in glutamatergic and GABAergic neurons (Luo et al., 2021). Our results show that baclofen-induced inhibition of DA overflow was not different across genotypes (**Fig. S1C**; Two-way ANOVA, F(1, 12)=0.008, *p*=0.93).

Together these observations suggest that Nrxns, while not required for DA release, play a role in regulating DA neurotransmission through regulation of DA reuptake and short-term plasticity in the vSTR. Our results suggest that Nrxns regulate DA release by means other than regulation of presynaptic GABA_B_ receptors.

### GABA release by dopamine neurons is increased in the ventral striatum of DAT::NrxnsKO mice

Whole-cell patch-clamp recordings in medium spiny neurons (MSNs) of the vSTR and dSTR in DAT::NrxnsWT, HET and KO littermates were performed to examine the role of Nrxns in synaptic GABA and glutamate release by DA neurons. A conditional AAV construct containing ChR2-EYFP was injected in the ventral mesencephalon and infected most of the neurons in the VTA and SNc of DAT-Cre::Nrxn mice (**Fig. 6A, 6B**). We isolated GABA-mediated synaptic currents (IPSC) pharmacologically and found a significant increase in the amplitude of IPSCs evoked by optical stimulation of DA neurons axons (oIPSCs) in the DAT::NrxnsKO slices (**Fig. 6C**; One-way ANOVA, F(2, 30) = 6.46, *p* = 0.005), with no change in electrically evoked GABA synaptic currents (eIPSC; **Fig. S3A**; One-way ANOVA, F(2, 35) = 1.46, *p* = 0.24). Interestingly, the frequency of spontaneous IPSCs (sIPSCs) was also significantly increased in the KO group (**Fig. 6D**; One-way ANOVA, F(2, 37) = 3.7, *p* = 0.03). oIPSCs were blocked by picrotoxin (50 µM), confirming their GABAergic nature (**Fig. S4A**). Furthermore, oIPSCs were blocked after treatment with the VMAT2 inhibitor reserpine (1 µM), in line with previous work showing that GABA release by DA neurons paradoxically requires VMAT2 (Tritsch et al., 2012; Tritsch et al., 2014) (**Fig. S4C**).

**Figure 6.**
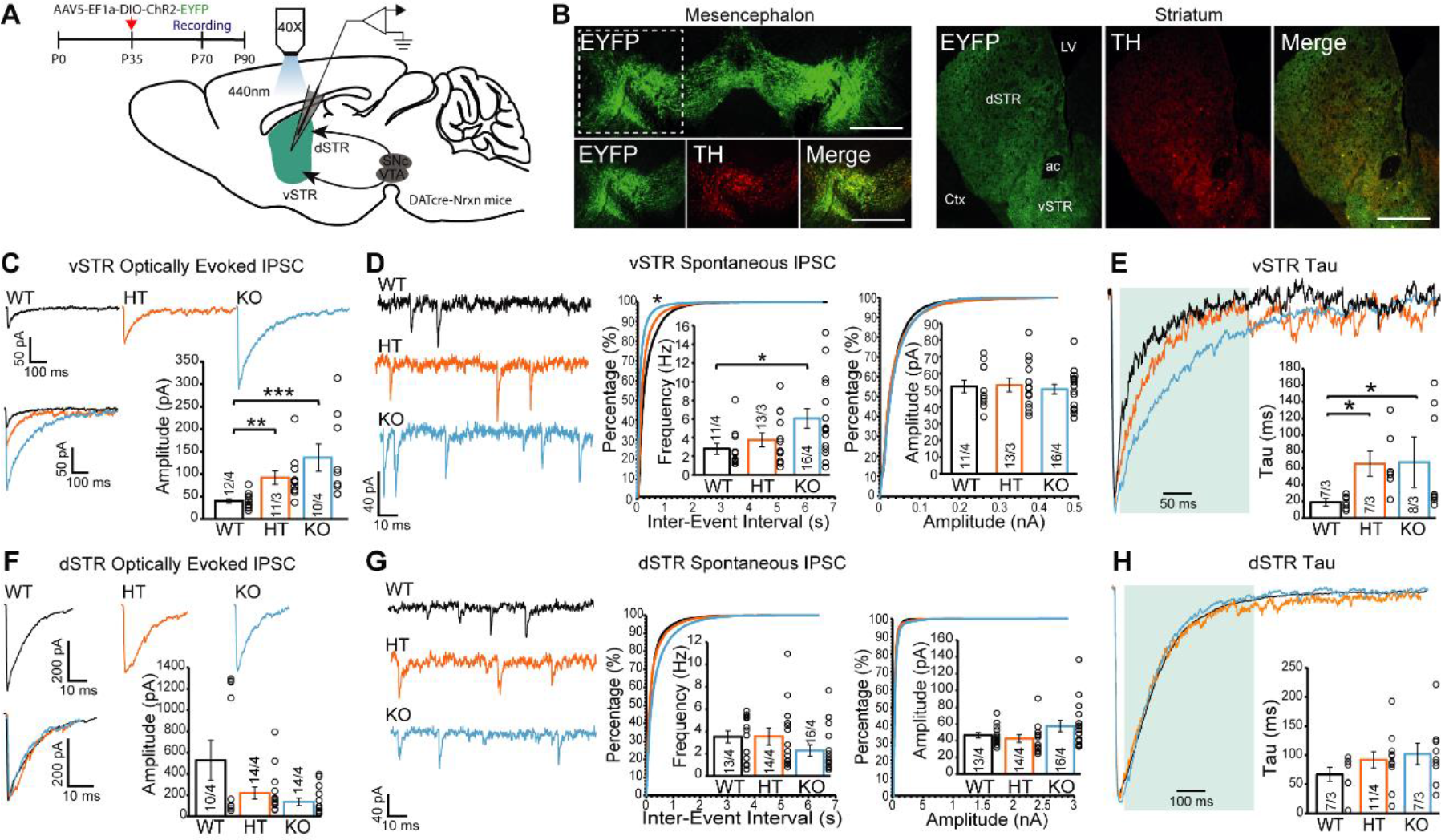
Neurexins regulate GABA release from dopaminergic neurons in the ventral striatum. **A-** Schematic representation of the experimental approach used for electrophysiological recordings; The green color represents the EYFP expression reporting ChR2 expression after AAV-transfection of the VTA. **B-** Photomicrographs illustrating the expression of EYFP-ChR2 in TH-positive (red) DA neurons located in the mesencephalon and their projections in the striatum. **C-** Representative traces (top) of IPSCs elicited by optical stimulation (o-IPSC), recorded in medium spiny neurons (MSNs) of acute ventral striatal slices from WT, HT, or KO mice. Summary graph of o-IPSC amplitude (bottom) shows a significant genotype effect (WT vs. HT: **p= 0.0017; WT vs. KO: ***p= 0.0001) (*Peak Amplitude:* WT = -40.02 ± 4.7 pA, HT = -91.99 ± 14.9 pA, KO = -136.32 ± 30.62, Mean ± SEM.). **D-** Representative traces (left) of spontaneous IPSCs (sIPSC) recorded in ventral striatal MSNs. Cumulative distribution plots of sIPSC inter-event intervals (middle, WT vs. KO: *p= 0.038) and amplitudes (right); insets: summary graphs of average sIPSC frequency (middle, WT vs. KO: *p=0.04) (*sIPSC frequency,* WT = 2.79 ± 0.61 Hz, HT = 3.72 ± 0.74 Hz, KO = 6.06 ± 1.04 Hz, Mean ± SEM) and amplitude (right). **E -** Representative traces and summary graph of o-IPSC decay time constant in ventral striatal MSNs, showing an increase in rate to return to baseline (oIPSC, decay tau: WT = 18.9 ± 2.8 ms, HT = 65.4 ± 13.5 ms. KO = 67.2 ± 21.9 ms. Mean ± SEM). **F -**Representative traces (left) of o-IPSCs recorded in MSNs of acute dorsal striatal slices from WT, HT or KO mice. The summary graph of o-IPSC amplitude (bottom) shows no genotype effect (Peak Amplitude: WT = -528.50 ± 187.31 pA, HT = -219.68 ± 54.87 pA, KO = -138.03 ± 34.57, Mean ± SEM). **G**- Representative traces (left) of sIPSCs recorded in dorsal striatal MSNs. Cumulative distribution plots of sIPSC inter-event intervals (middle) and amplitudes (right); insets: summary graphs of average sIPSC frequency (middle) and amplitude (right). Summary graphs show no genotype effect **H-** Representative traces and summary graph of o-IPSC decay time in dorsal striatal MSNs shows no change in tau for both HT and KO compared to the WT group. Data in summary graphs are presented as mean ± SEM; statistical comparisons were performed with One-Way ANOVA (*P<0.05; **P<0.01; ***P<0.001; non-significant comparisons are not identified). Tukey’s correction was used for all multiple comparisons. The Kruskal-Wallis non-parametric test was used to compare cumulative distributions in **D** and **G**. Numbers in bars indicate the number of cells/mice. Each open circle in the summary graphs represents the averaged recording of each cell.

Closer analysis of the kinetics of GABA-mediated oIPSCs in the vSTR revealed a significant increase of oIPSC decay time (**Fig. 6E, S5A-D**; One-way ANOVA, F(2, 21) = 4.43, *p* = 0.02) and decay time constant (**Fig. S5**; One-way ANOVA, F(2, 19) = 5.55, *p* = 0.013) in both DAT::NrxnsKO and DAT::NrxnsHET mice. These findings suggest that neurexin deletion in vSTR DAergic terminals enhances GABA release from DA terminals and prolongs GABAergic activity in MSNs.

As our manipulation affected the entire mesolimbic pathway, we also performed parallel optical stimulation and recordings in the dorsal striatum. We observed no statistically significant changes in oIPSC amplitude in DAT::NrxnsKO mice in the dSTR (**Fig. 6F**; Kruskal-Wallis Test, H(3) = 3.4, *p* = 0.18). Spontaneous and electrically-evoked GABA IPSCs were also unaltered (**Fig. 6G**, **Fig. S3C and S3D**), as was oIPSC decay time constant (**Fig. 6H** and **S5E-H**; One-way ANOVA, F(2, 24) = 1.2, *p* = 0.32).

Together these results suggest that Nrxns act as regulators of GABA co-transmission by DA neurons in a region-specific manner. Further work will be required to identify the origin of this selectivity.

### Optically evoked excitatory transmission is unaltered in the striatum of DAT::NrxnsKO mice

We also examined glutamate release by DA neurons in the vSTR and dSTR. We did not detect genotype differences in optically-evoked EPSCs (oEPSCs) (**Fig. 7A**; One-way ANOVA, F(2, 30) = 1.01, *p* = 0.37**, 7B**; One-way ANOVA, F(2, 20) = 0.8, *p* = 0.46). The frequency and amplitude of spontaneous EPSCs (sEPSCs) was also unchanged (**Fig. 7C, 7D**). The paired-pulse ratio and input-output relationship of electrically-evoked IPSCs were similarly unchanged (**Fig. S6**). These results suggest that Nrxns do not have a major role in regulating glutamate release from DA neurons. However, we detected a significant increase of the amplitude of EPSCs evoked by local electrical stimulation (eEPSC) in the ventral (**Fig. S3B**; One-way ANOVA, F(2, 30) = 4.8, *p* = 0.01) but not the dorsal (**Fig. S3D**; One-way ANOVA, F(2, 29) = 0.02, *p* = 0.98) striatum. This observed increase in eEPSC amplitude opens the possibility that deletion of Nrxns impacts the development of DAergic projections to non-striatal regions that project to vSTR.

**Figure 7.**
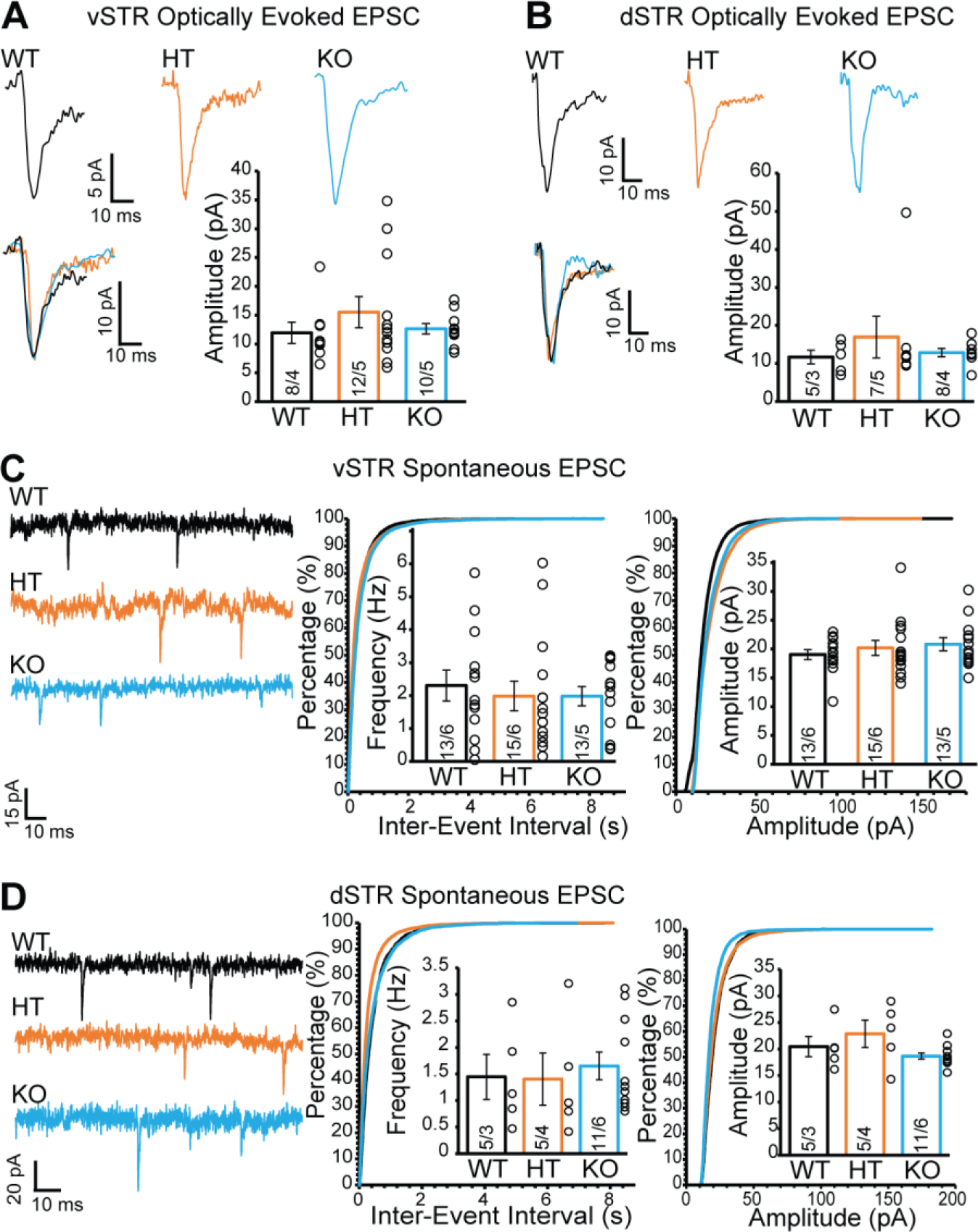
Loss of neurexins does not influence glutamate release from dopamine neurons in the ventral and dorsal striatum. **A**- Representative traces (top) of EPSCs elicited by optical stimulation (o-EPSCs) and recorded in ventral striatal MSNs. Summary graph of o-EPSC amplitudes (bottom) shows no genotype effect. **B**- Representative traces (top) of o-EPSCs recorded in dorsal striatal MSNs. Summary graph of o-EPSC amplitudes (bottom) shows no genotype effect. **C**- Representative traces (left) of spontaneous EPSCs (sEPSC) recorded in ventral striatal MSNs. Cumulative distribution plots of sEPSC inter-event intervals (middle) and amplitudes (right); insets: summary graphs of average sEPSC frequency (middle) and amplitude (right). **D**- Representative traces (left) of spontaneous EPSC (sEPSC) recorded in dorsal striatal MSNs. Cumulative distribution plots of sEPSC inter-event intervals (middle) and amplitudes (right); insets: summary graphs of average sEPSC frequency (middle) and amplitude (right). Data in summary graphs are provided as mean ± SEM. Statistical comparisons were performed with One-Way ANOVA (*P<0.05; **P<0.01; ***P<0.001; non-significant comparisons are not labeled). Tukey’s correction was used for all multiple comparisons.

### Altered GABA uptake and VMAT2 suggest a possible mechanism underlying increased GABA release in Nrxns KO mice

The region-specific increase of GABA release from DA axons in DAT::NrxnsKO mice, identified in our optogenetic experiments, could result from different mechanisms. One possibility is that loss of Nrxns induced differential adaptations in the expression of some of the postsynaptic partners of Nrxns, including NLs. However, using qRT-PCR, we did not detect major changes in expression of these genes in microdissected vSTR or dSTR, except for a small decrease in collybistin and LRRTM3 mRNA in DAT::NrxnsKO mice (unpaired t-test, *p*=0.021 and *p*=0.014, respectively) (**Fig. S7A** and **S7B**).

While DA neurons have the capacity to co-release GABA, it is well established that they do not synthesize it using glutamic acid decarboxylase and do not express the vesicular GABA transporter (Tritsch et al., 2012; Tritsch et al., 2014). Instead, they have been shown to use plasma membrane transporters to uptake GABA from the extracellular medium and VMAT2 to package it into synaptic vesicles (Tritsch et al., 2012; Tritsch et al., 2014). We therefore used a GABA uptake assay using primary DA neurons and quantified VMAT2 levels in the striatum.

GABA uptake was estimated using cultures prepared from DA neurons co-cultured with vSTR or dSTR neurons. Neurons were incubated with GABA (100µM) for 2h, rapidly washed and fixed before quantification of GABA immunoreactivity in these neurons, evaluating the proportion of TH signal in DA axons covered by GABA immunoreactivity in comparison to a control group treated with H_2_O (**Fig. 8A**). This treatment induced a robust increase in GABA immunoreactivity in DA neuron axons (**Fig. 8B** to **8D**). Interestingly, SNc DA neurons globally showed a larger GABA uptake compared to VTA DA neurons (**Fig. 8E**; two-way ANOVA, main effect of region, F(1, 76)=53.35, *p<*0.001). Furthermore, we observed a global increase of GABA uptake in DAT::NrxnsKO DA neurons compared to DAT::NrxnsWT DA neurons (two-way ANOVA, main effect of genotype, F(1, 76)=6.25, *p*=0.014).

**Figure 8.**
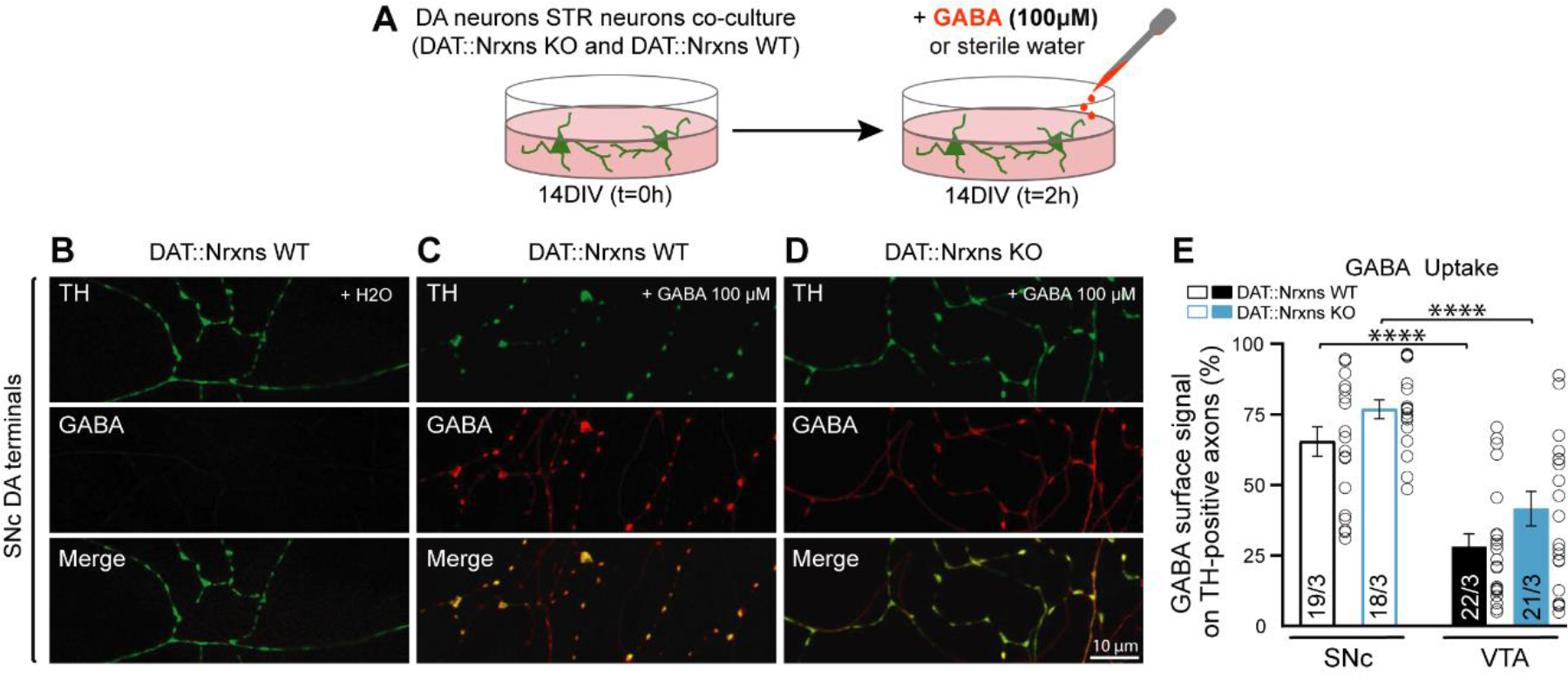
GABA uptake from cultured DA neurons is unchanged after conditional deletion of all neurexins. **A** - Schematic representation of the experimental procedure for the GABA uptake assay in VTA-vSTR co-cultures or in SNc-dSTR co-cultures. **B - D-** Immunocytochemistry of SNc DA neurons from Nrxn123 WT (B and C) and DAT::NrxnsKO mice (D) for tyrosine hydroxylase (TH, green) and gamma-aminobutyric acid (GABA, red). Experiments on VTA-vSTR co-cultures are not illustrated. **E** - Summary graph representing the quantification of GABA immunoreactivity signal surface in TH-positive axons for VTA and SNc DA neurons from DAT::NrxnsWT and KO cultures. N = 18-22 axonal fields from 3 different neuronal co-cultures. The number of observations represents the number of fields from TH-positive neurons examined. For all analyses, plots represent the mean ± SEM. Statistical analyses were carried out by 2-way ANOVAs followed by Tukey’s multiple comparison test. (*p < 0.05; **p < 0.01; ***p < 0.001; ****p < 0.0001).

We next examined the level of VMAT2, TH and DAT in DA neuron axons in the striatum of DAT::NrxnsKO mice compared to WT controls. A first global quantification of protein levels by Western Blot (**Fig. S8A and S8E**), revealed a lack of overall change in the levels of TH and VMAT2 in DAT::NrxnsKO mice, in both vSTR and dSTR tissues (**Fig. S8B, S8C, S8F and S8G**). However, DAT levels were significantly reduced in the vSTR (**Fig. S8A** and **S7D**; unpaired t-test, t_10_=2.79; *p*=0.019). We also observed a tendency for a similar decrease in DAT in the dSTR (**Fig. S8E** and **S8H;** unpaired t-test, t_10_=1.93; *p*=0.081).

To gain more region-specific insight in the levels of these proteins, we measured the intensity and surface area for these markers in a series of 3 striatal brain sections ranging from bregma +0.74 to bregma -0.82 mm, with a total of 7 different regions in each hemisphere (**Fig. 9A**). In this experiment, all sub-regions were compared and analyzed separately. First, TH signal intensity (**Fig. 9B, 9C and 9F**) and TH surface area (**Fig. 9G**) were unchanged in both dSTR and vSTR. For VMAT2 immunoreactivity (**Fig. 9B and 9C**), surface area was increased in both the vSTR and in the rostral part of the dSTR for KO mice (**Fig. 9I;** vSTR, unpaired t-test, Welch’s correction, *p*=0.045; dSTR3, unpaired t-test, Welch’s correction, *p*=0.045), while intensity was unchanged (**Fig. 9H**). In contrast, DAT immunostaining showed a decrease of signal intensity in the vSTR (**Fig. 9D** and **9J**; vSTR, unpaired t-test, *p*=0.039) and in the rostral and caudal parts of the dSTR of DAT::NrxnsKO mice (**Fig. 9D, 9E** and **9J**; dSTR1, unpaired t-test, *p*=0.025; dSTR3, unpaired t-test, Welch’s correction, *p*=0.024). The DAT surface area was also significantly decreased in the vSTR and in the caudal part of the dSTR (**Fig. 9K**; vSTR, unpaired t-test, *p*=0.033; dSTR3, unpaired t-test, Welch’s correction, *p*=0.030).

**Figure 9.**
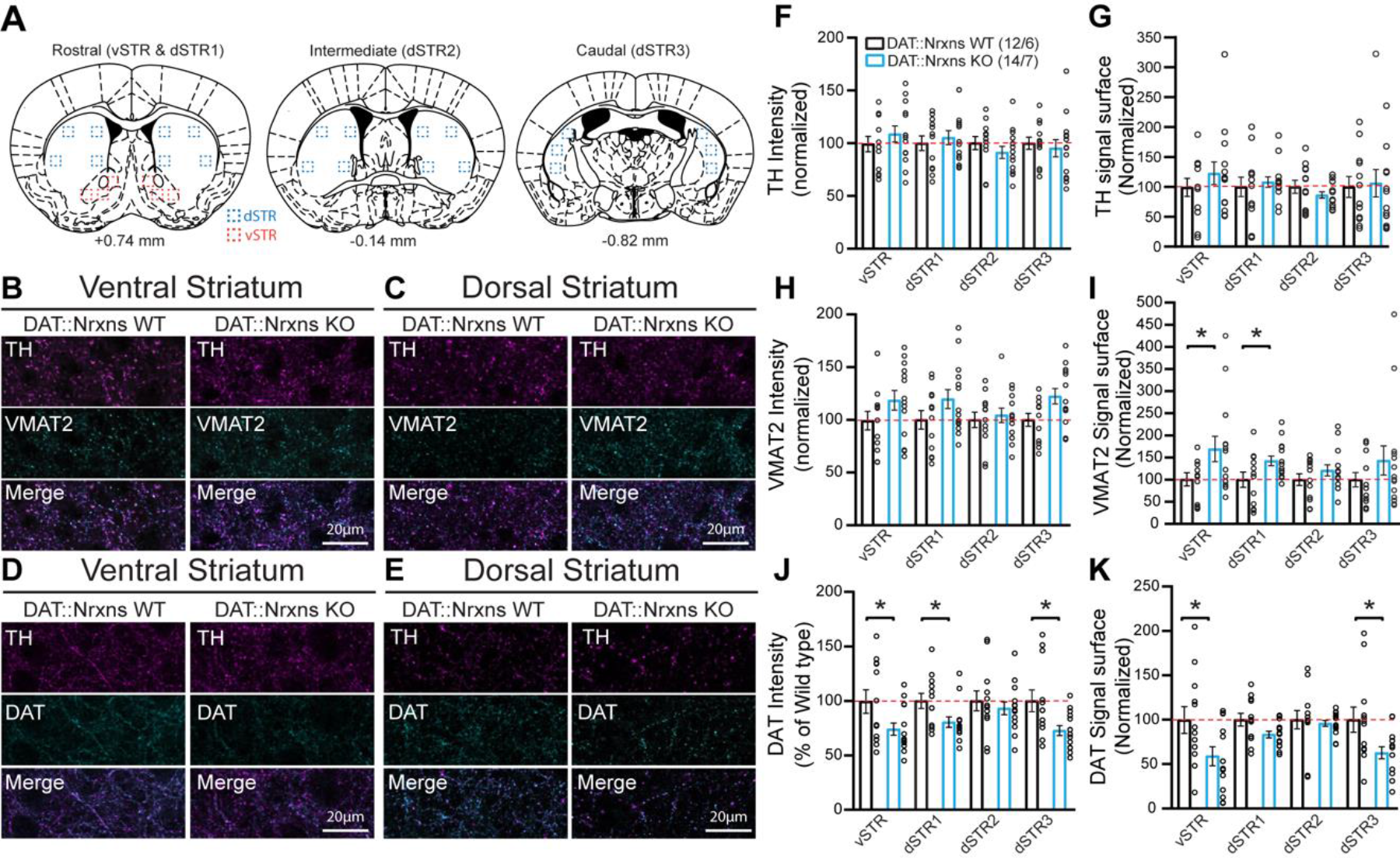
Increased VMAT2 but decreased DAT expression in DA axon terminals lacking neurexins. **A**- Schematic representation of striatal slices used for immunohistochemistry characterization. Surface and intensity for each signal were measured in a series of 3 different striatal slices ranging from bregma +0.74 to bregma -0.82 mm, with a total of 14 different spots for each hemisphere. **B and C** - Immunohistochemistry of ventral (**B**) and dorsal (**C**) striatal slices from 8-week-old DAT::NrxnsKO and DAT::NrxnsWT mice (60X confocal) using tyrosine hydroxylase (TH, purple) and vesicular monoamine transporter 2 (VMAT2, blue) antibodies. **D and E** - Immunohistochemistry of ventral (**B**) and dorsal (**C**) striatal slices from DAT::NrxnsKO and DAT::NrxnsWT mice using TH (purple) and the dopamine transporter (DAT, blue) antibodies. **F to K** - Quantification of signal surface (% of WT) for TH, VMAT2 and DAT in the different striatal regions examined: “vSTR, dSTR1, dSTR2 and dSTR3” (DAT::NrxnsKO = 14 hemispheres/7 mice; DAT::NrxnsWT = 12 hemispheres/6 mice). TH signal intensity: vSTR= 108.7 ± 7.48%; dSTR1= 105.2 ± 6.65%; dSTR2= 91.18 ± 5.71%; dSTR3= 95.26 ± 8.05% of control. TH surface area: vSTR= 122.6 ± 19.07%; dSTR1=107.9 ± 8.62%; dSTR2= 86.38 ± 5.42%; dSTR3= 106.20 ± 22.82% of control. VMAT2 intensity: vSTR= 118.4 ± 9.30%; dSTR1= 119.7 ± 8.96%; dSTR2= 104.3 ± 6.91%; dSTR3= 122.3 ± 7.24% of control. VMAT2 surface area: vSTR=169.3 ± 28.42% dSTR1=142.4 ± 11.14% dSTR2=120.8 ± 12.73% dSTR3=143.3 ± 32.98%. DAT signal intensity: vSTR=74.07 ± 20.98%; dSTR1=80.50 ± 17.77%; dSTR2=93.17 ± 21.96%; dSTR3=72.80 ± 17.08%. DAT surface area: vSTR=58.92 ± 10.72%; dSTR1=60.80 ± 3.87%; dSTR=295.98 ± 3.38%; dSTR3=62.78 ± 6.71%. Statistical analysis was carried out by unpaired t-test for each substructure. Error bars represent ± S.E.M. (ns, non-significant; *, p < 0.05; **, p < 0.01; ***, p < 0.001; ****, p < 0.0001).

Together, our observation of increased GABA uptake and increased VMAT2 in the vSTR could represent an explanation for the observed increase in GABA release by DA neuron axons in this part of the striatum.

## Discussion

Since the initial discovery of Nrxns (Ushkaryov et al., 1992), multiple studies have explored the roles of these proteins in synapse formation, function, maintenance, and plasticity. Most of these studies have been conducted on glutamatergic or GABAergic neurons, with no evaluation of their role in modulatory neurons such as DA, serotonin, norepinephrine, or acetylcholine neurons, whose connectivity is strikingly different and markedly less synaptic (Ducrot et al., 2021). We expected that new insights could be gained by studying the role of these trans-synaptic proteins in modulatory neurons. In the present study, we utilized the Cre-lox system by combining the triple conditional Nrxn mouse line (Chen et al., 2017) with a DAT-Cre mouse line to selectively delete Nrxns in DA neurons and examine the impact of this deletion using a combination of primary neuronal cultures, electron microscopy, behavioral assessments, patch-clamp recordings of striatal MSNs, fast scan cyclic voltammetry, and immunohistochemistry. We found that while loss of Nrxns does not markedly impair the basic development and axonal ultrastructure of DA neuron terminals *in vitro* and *in vivo*, it does lead to region-specific changes in their axonal connectivity. We also found that loss of Nrxns is associated with impaired DA neurotransmission in the brain of adult mice, as revealed by a reduced rate of DA reuptake after electrically-evoked DA release and with impaired amphetamine-induced locomotion. Patch-clamp recordings of GABA and glutamate release by DA neuron axons also revealed an unexpected increase of GABA co-release. Together these findings suggest that, although Nrxns are not critical for many aspects of the basic axonal development and synapse formation of DA neurons, they act as key regulators of GABA and DA signaling in these neurons.

### Nrxns are not required for the basic morphological development of DA neurons but regulate their connectivity

Nrxns have been previously suggested to contribute to the development of synapses (Missler et al., 2003; Li et al., 2007; Etherton et al., 2009; Aoto et al., 2015; Chen et al., 2017). In a first series of experiments, we examined whether loss of Nrxns impairs the morphological development of DA neurons. Our results did not identify major changes in the global axonal development of these neurons. However, we found that the proportion of bassoon-positive terminals, established by SNc DA neurons, was reduced after loss of Nrxns. Furthermore, the proportion of these terminals contacting PSD95 postsynaptic domains was also reduced, suggesting that these proteins play a role in regulating the connectivity of DAergic terminals, and the formation of active zones. These findings are in line with recent results suggesting a possible role of Nrxns in the organizational features of the active zone (Luo et al., 2020). How Nrxns organize active zone assembly is presently unknown, but a previous study suggested that these proteins can interact indirectly with active zone proteins through calcium/calmodulin-dependent serine protein kinase 3 and membrane-associated guanylate kinase 2 (CASK) (Hata et al., 1996). Further *in vivo* experiments will be needed to evaluate this possibility more directly using transmission electron microscopy or other types of high-resolution microscopy approaches.

Experiments performed *in vitro* allowed us to more closely examine the propensity of DA terminals to interact with GABA and glutamate postsynaptic domains, especially considering the capacity of these neurons to co-release glutamate and GABA in a subset of their synaptic terminals (Sulzer et al., 1998; Dal Bo et al., 2008; Descarries et al., 2008; Stuber et al., 2010; Tritsch et al., 2012). We hypothesized that Nrxns play a critical role in the establishment of excitatory and inhibitory synapses by DA neurons. Quantification of the proportion of TH-positive terminals overlapping with the glutamatergic postsynaptic marker PSD95 revealed a decrease in the proportion of SNc DA neuron terminals contacting a PSD95 or gephyrin postsynaptic domain in cultures prepared from DAT::NrxnsKO mice. However, results obtained with transmission electron microscopy in striatal sections did not reveal any notable differences in the ultrastructure of DA terminals in these mice when compared to wild-type controls. However, these experiments did not distinguish between DA neuron terminals containing vesicular glutamate transporters and those only containing DA markers. Further immuno-EM experiments would be required to examine VGLUT2-positive DA neuron terminals more specifically. These data suggest that although Nrnxs are not required for the basic structure of DA neuron terminals, they can regulate DA neuron connectivity in a region-specific manner. Our observations of region-specific alterations concur with results from the first *in vivo* study on the conditional triple Nrxn deletion, published by Chen and colleagues in 2017 (Chen et al., 2017), revealing that removal of Nrxns resulted in a significant decrease in the number of parvalbumin-positive synapses in the cortex and of excitatory synapses formed by the climbing fibers innervating purkinje neurons in the cerebellum. Our observation of an absence of major structural changes in the DA neuron terminals after deletion of Nrxns is also in keeping with previous reports obtained with the single, double or triple deletion of Nrxn in glutamatergic or GABAergic neurons (Missler et al., 2003; Chen et al., 2017). Another recent study using the triple Nrxn mice also reported a lack of impact of Nrxn deletion on synapse formation at the calyx of Held (Luo et al., 2020). Taken together with our results, we similarly conclude that Nrxns play a synapse-specific regulatory role in synapse formation and maintenance, but do not act as direct drivers of synapse formation.

### Dopamine transporter expression and function are altered in DAT::NrxnsKO mice

To obtain direct functional insight into the roles of Nrxns in DAergic neurotransmission, we performed FSCV recordings in the vSTR and dSTR. Our data revealed that peak electrically-evoked DA overflow was not reduced in the absence of Nrxns, suggesting that Nrxn123 do not play an obligatory role in the DA release mechanism and the initial formation and function of DA neuron varicosities, in line with our *in vitro* results. The reduction in peak DA overflow in DAT::NrxnsHET mice is puzzling and remains presently unexplained. One possibility is that deletion of a single Nrxn123 allele was insufficient to prevent the occurrence of possible compensatory mechanisms induced in full knockouts. Although peak DA release was not impaired in the absence of Nrxns, we found a systematic impairment in DA reuptake in both DAT::NrxnsKO and DAT::NrxnsHET mice compared to DAT::NrxnsWT mice. This finding of impaired DAT function is particularly intriguing in the context of our observation that DAT::NrxnsKO mice show a robust impairment in amphetamine-induced locomotion. Amphetamine is well known to increase extracellular DA levels by impairing vesicular storage of DA and inducing reverse transport through the DAT. It is thus possible that Nrxns regulate the stability and function of the DAT in DA neuron axon terminals, perhaps through DAT’s PDZ domain, and that in the absence of Nrxns, DAT function and positioning in terminals is impaired (Sorensen et al., 2021).

### Regulatory role of Nrxns in GABA release by dopamine neurons

One of the most intriguing observations in the present study is the region-specific increase of evoked GABA release from VTA DA neuron terminals in the vSTR. The origin of this selectivity is presently unclear but could arise from several possibilities including, differential expression of Nrxn splice variants or their postsynaptic ligands and selective binding affinity in the dorsal or ventral striatum. Multiple studies over the past decade reported similar conclusions, where different Nrxn isoforms were proposed to regulate various aspects of synapse organization and function (Ullrich et al., 1995). Further studies would be needed to analyze the function of specific splice variants in each region. It is known that Nrxn splice variants are differentially expressed in different brain regions, and this determines the preference for different postsynaptic binding partners. A possible hypothesis to explain our observations is that one or more Nrxns act as a repressor of GABA co-transmission and thus regulate the excitatory/inhibitory neurotransmission balance of the axonal domain of DA neurons. Indeed, previous work has shown that Nrxns physically and functionally interact with GABA_A_ receptors and that overexpression of Nrxns decreases inhibitory but not excitatory synaptic strength (Zhang et al., 2010). These results are consistent with our observations showing that after removal of all Nrxns, GABAergic synaptic events evoked by stimulation of DA neuron terminals are increased. Further work, including rescue experiments, would be needed to test this hypothesis directly and determine which Nrxn is involved in this mechanism.

Interpretation of some of our findings needs to be made considering that the DAT promoter becomes active during the late embryonic period. Embryonic deletion of Nrxns in DA neurons could thus have altered the development of these neurons and of target regions. Notably, our observation of an increase in electrically evoked EPSCs and sIPSCs in the vSTR of KO mice could potentially result from alterations in DA neuron projections to non-striatal regions such as the cortex.

The regional specificity of our data raises intriguing questions regarding functional differences between the mesolimbic and nigrostriatal projections. We observed an increase in GABA-mediated oIPSCs only in the vSTR. Furthermore, basal GABA uptake was higher in SNc than VTA DA neurons and baseline oIPSC amplitude was higher in the dSTR than the vSTR. Together, these observations suggest that there may be intrinsic differences in the structure and function of GABA release sites in these two circuits.

The role of Nrxns in regulating glutamate synapses established by DA neuron axons is less clear. Although we detected a decrease in the propensity of DA neuron axonal varicosities to contact PSD95 postsynaptic domains, we did not detect a decrease in optically-evoked glutamate-mediated EPSCs in striatal neurons in slices prepared from DAT::NrxnsKO mice. This dichotomy suggests that although structural changes may have happened at these glutamatergic terminals, this did not translate into functional changes. These findings highlight the complexity and the diversity of the role of Nrxns at synapses, in line with much recent work (Chen et al., 2017; Luo et al., 2020).

It would be interesting to complement our work by recording separately from striatal MSNs of the direct and indirect basal ganglia pathway (Gerfen and Surmeier, 2011), as some of the heterogeneity in our results on GABA and glutamate synaptic events may derive from differences in the roles of Nrxns in these two pathways. Previous work has shown that DA regulates tonic inhibition in striatal MSNs, and this regulation differs between D1 and D2 MSNs (Janssen et al., 2009). Given our results showing changes in GABA release from DA terminals, investigating tonic inhibition of D1 and D2 MSNs within the context of Nrxn deletion would be of interest for future work.

Mechanistically, the increase in GABA IPSCs we detected in DAT::NrxnsKO mice most likely resulted from presynaptic changes. We detected an increase in GABA uptake by cultured DA neurons and an increase in VMAT2 levels, detected in striatal sections. Together these two mechanisms could lead to increased GABA uptake by DA neuron terminals and increased GABA vesicular packaging through a VMAT2-dependent process. We also confirmed that oIPSCs in the vSTR were blocked almost entirely by reserpine, a VMAT2 inhibitor. Importantly, the increase in oIPSC amplitude in the vSTR was not accompanied by changes in eIPSC amplitude or sIPSC amplitude. This further supports the idea that the changes observed in GABA-mediated currents resulted specifically from an increase in presynaptic GABA release from DA neurons. However, the prolongation of optically-evoked IPSCs that we also detected in slices prepared from DAT::NrxnsKO mice points to a prolongation of GABA_A_ receptor opening. Further experiments will be required to identify the pre- or postsynaptic origin of this effect.

Alterations in DA neuron connectivity following Nrxn deletion may also regulate the intrinsic vulnerability of DA neurons, as suggested by our observation of reduced survival of DA neurons in primary cultures established from DAT::NrxnsKO mice. However, we did not observe a decrease in TH signal in the striatum of these mice *in vivo*.

Together, our findings shed new light on the role of these cell-adhesion molecules in DA neuron connectivity. We conclude that Nrxns are dispensable for the initial establishment of axon terminals and synapses by DA neurons but play a role in regulating both DAT function and GABA release by DA neurons. Only a small subset of axon terminals established by DA neurons adopt a synaptic configuration. We conclude that the formation of such synaptic contacts must be regulated by other transsynaptic proteins. Potential candidates include proteins from the leucocyte antigen receptor – protein tyrosine-phosphatases (LAR-PTP) family. As we observed a decrease in bassoon-positive terminals established by DA neurons in the absence of Nrxns, it remains possible that these trans-synaptic proteins play a role in regulating active zone assembly in a subset of terminals established by DA neurons. Further experiments will be needed to determine if this assembly is subject to control by presynaptic signals or if it is controlled indirectly by trans-synaptic signaling via postsynaptic signals.

## STAR methods

### Animals

All procedures involving animals and their care were conducted in accordance with the Guide to care and use of Experimental Animals of the Canadian Council on Animal Care. The experimental protocols were approved by the animal ethics committees of the Université de Montréal (CDEA). Housing was at a constant temperature (21°C) and humidity (60%), under a fixed 12h light/dark cycle with food and water available ad libitum.

### Generation of triple Neurexins cKO mice in DA neurons

All experiments were performed using mice generated by crossing DAT-IRES-Cre transgenic mice (Jackson Labs, B6.SJL-Slc6a3tm1.1 (Cre)Bkmn/J, strain 006660) with Nrxn123loxP mice [for details see (Chen et al., 2017)]. Briefly, conditional knock-out (cKO) mice were produced as a result of CRE recombinase driving a selective excision of Nrxn-1, Nrxn-2 and Nrxn-3 genes in DA neurons and giving three different genotypes: Nrxn123 KO, Nrxn123 HET and Nrxn123 WT mice. The Nrxn123flox/flox mice were on a mixed Cd1/BL6 genetic background. The DAT-IRES-Cre mice were on a C57BL/6J genetic background. Except for culture experiments, only males were used.

### Genotyping

Mice were genotyped with a KAPA2G Fast HotStart DNA Polymerase kit (Kapa Biosystem). The following primers were used: DAT-IRES-Cre: Common 5’ TGGCTGTTGTGTAAAGTGG3’, wild-type reverse 5’GGACAGGGACATGGTTGACT 3’ and knock-out reverse 5’-CCAAAAGACGGCAATATGGT-3’, Nrxn-1 5’-GTAGCCTGTTTACTGCAGTTCATT-3’ and 5’-CAAGCACAGGATGTAATGGCCTT-3’, Nrxn-2 5’-CAGGGTAGGGTGTGGAATGAG-3’ and 5’-GTTGAGCCTCACATCCCATTT-3’, Nxn3 5’-CCACACTTACTTCTGTGGATTGC-3’ and 5’-CGTGGGGTATTTACGGATGAG-3’.

### Primary neuronal co-cultures

For all experiments, postnatal day 0-3 (P0-P3) mice were cryoanesthetized, decapitated and used for co-cultures as previously described (Fasano et al., 2008). Primary VTA or SNc DA neurons were obtained from Nrxn123 KO or Nrxn123 WT pups and co-cultured with ventral striatum and dorsal striatum neurons, respectively from Nrxn123 KO or Nrxn123 WT pups. Neurons were seeded on a monolayer of cortical astrocytes grown on collagen/poly-L-lysine-coated glass coverslips. All cultures were incubated at 37°C in 5% CO2 and maintained in 2/3 of Neurobasal medium, enriched with 1% penicillin/streptomycin, 1% Glutamax, 2% B-27 supplement and 5% fetal bovine serum (Invitrogen) plus 1/3 of minimum essential medium enriched with 1% penicillin/streptomycin, 1% Glutamax, 20mM glucose, 1mM sodium pyruvate and 100 µl of MITO+ serum extender. All primary neuronal co-cultures were used at 14DIV.

### Immunocytochemistry on cell cultures

Cultures were fixed at 14-DIV with 4% paraformaldehyde (PFA; in PBS, pH-7.4), permeabilized with 0,1% triton X-100 during 20-min, and nonspecific binding sites were blocked with 10% bovine serum albumin during 10-min. Primary antibodies were: mouse anti-tyrosine hydroxylase (TH) (1:2000, Sigma), rabbit anti-TH (1:2000, Chemicon), rabbit anti-synaptotagmin 1 (Syt1) (1:1000, Synaptic Systems) and rabbit anti-vesicular monoamine transporter 2 (VMAT2) (1:1000, gift of Dr. Gary Miller, Colombia University). To improve the immunoreactivity of the synaptic markers PSD95, gephyrin and bassoon, a set of cultures were fixed with 4% PFA together with 4% sucrose. For these experiments, the primary antibodies were mouse anti-PSD95 (1:1000 Pierce antibody), mouse anti-gephyrin (1:1000, Synaptic Systems) and guinea pig anti-bassoon (1:1000, Synaptic Systems). These were subsequently detected using Alexa Fluor-488-conjugated, Alexa Fluor-546-conjugated, Alexa Fluor-568-conjugated or Alexa Fluor-647-conjugated secondary antibodies (1:500, Invitrogen).

### Transmission Electron Microscopy (TEM)

Following i.p. injection of ketamine (100 mg/Kg) and xylazine (10 mg/Kg), P70 mice were transcardially perfused with 50 ml of ice-cold sodium phosphate buffer saline (PBS; 0.1M; pH 7.4) followed by 150 mL of a mix composed of 4% paraformaldehyde and 0.1% glutaraldehyde diluted in PBS. Dissected brains were extracted and post-fixed overnight in 4% PFA at 4°C. Mouse brains were cut with a vibratome (model VT1200 S; Leica, Germany) into 50 µm-thick transverse sections. For TH immunostaining, 50 µm-thick sections taken through the striatum (1.18 mm and 1.34 mm from bregma, according to the mouse brain atlas of Franklin & Paxinos 1^st^ edition) were rinsed with PBS and pre-incubated for 1h in a solution containing 2% normal goat serum and 0.5% gelatin diluted in PBS. Sections were then incubated overnight with a rabbit primary TH antibody (Millipore, catalogue no. AB152, 1/1000). Sections were rinsed and incubated during 2h with a goat anti-rabbit secondary antibody (1/500) and directly coupled to a peroxidase (Jackson, catalogue no. 111-035-003). The peroxidase activity was revealed by incubating sections for 5 min in a 0.025 % solution of 3,3’ diaminobenzidine tetrahydrochloride (DAB; Sigma-Aldrich, catalogue no. D5637) diluted in Tris-saline buffer (TBS; 50 mM; pH 7.4), to which 0.005% of H₂O₂ was added. The reaction was stopped by several washes in TBS followed by phosphate buffer (PB; 50 mM; pH 7.4). At room temperature, sections were washed 3 times in ddH_2_O and incubated for 1h in a solution composed of 1.5% potassium ferrocyanide and 2% osmium tetroxide (EMS) diluted in ddH_2_O. After 3 rinses in ddH_2_O, sections were incubated for 20 min in a filtered solution of 1% thiocarbohydrazide (Ted Pella) diluted in ddH_2_O. Sections were then rinsed 3 times and incubated in 2% osmium tetroxide. After rinses in ddH_2_O, sections were dehydrated in graded ethanol and propylene oxide and flat-embedded in Durcupan (catalogue no.44611-14; Fluka, Buchs, Switzerland) for 72h at 60°C. Trapezoidal blocs of tissue from the ventral striatum were cut from the resin flat-embedded TH-immunostained sections. Each quadrangular pieces of tissue were glued on the tip of resin blocks and cut into 80 nm ultrathin sections with an ultramicrotome (model EM UC7, Leica). Ultrathin sections were collected on bare 150-mesh copper grids and examined under a transmission electron microscope (Tecnai 12; Philips Electronic, Amsterdam, Netherlands) at 100 kV. Profiles of axon varicosities were readily identified as such by their diameter (larger than 0.25 µm) and their synaptic vesicles content. Using an integrated digital camera (MegaView II; Olympus, Münster, Germany), TH immunopositive axon varicosities were imaged randomly, at a working magnification of 9,000X, by acquiring an image of every such profile encountered, until 50 or more showing a full contour and distinct content were available for analysis, in each mouse.

### Stereotaxic virus injections

Nrxn123 KO, Nrxn123 HET and Nrxn123 WT mice (P30-45) were anesthetized with isoflurane (Aerrane; Baxter, Deerfield, IL, USA) and fixed on a stereotaxic frame (Stoelting, Wood Dale, IL, USA). Fur on top of the head was trimmed, and the surgical area was disinfected with iodine alcohol. Throughout the entire procedure, eye gel (Lubrital, CDMV, Canada) was applied to the eyes and a heat pad was placed under the animal to keep it warm. Next, bupivacaine (5-mg/ml and 2-mg/Kg, Marcaine; Hospira, Lake Forest, IL, USA) was subcutaneously injected at the surgical site, an incision of about 1-cm made with a scalpel blade and the cranium was exposed. Using a dental burr, one hole of 1-mm diameter was drilled above the site of injection [AP (anterior-posterior; ML (medial-lateral); DV (dorsal ventral), from bregma]. The following injection coordinates were used: SNc/VTA [AP -3.0 mm; ML ± 0.9 mm; DV -4.3 mm]. Borosilicate pipettes were pulled using a Sutter Instrument, P-2000 puller, coupled to a 10 ul Hamilton syringe (Hamilton, 701 RN) using a RN adaptor (Hamilton, 55750-01) and the whole setup was filled with mineral oil. Using a Quintessential Stereotaxic Injector (Stoelting), solutions to be injected were pulled up in the glass pipette. For expression of ChR2 in DA neurons, 0.8-µl (VTA/SNc) of sterile NaCl containing 1.3x10^12^ viral particles/mL of AAV5-EF1a-DIO-hChR2(H134R)-EYFP (UNC GTC Vector Core, NC, USA) was injected bilaterally. After each injection, the pipette was left in place for 5-min to allow diffusion and then slowly withdrawn. A second batch of mice were injected twice bilaterally [AP -2.8 mm; ML ± 0.9 mm; DV -4.3 mm] and [AP -3.2 mm; ML ± 1.5 mm; DV -4.2 mm] with a total of four 0.5-uL injections of the same viral preparation. This was done to improve the infection rate. Finally, the scalp skin was sutured and a subcutaneous injection of the anti-inflammatory drug carprofen (Rimadyl, 50-mg/mL at 5 mg/kg) was given. Animals recovered in their home cage and were closely monitored for 72h. A second dose of carprofen (5-mg/kg) was given if the mice showed evidence of pain.

### Electrophysiology and optogenetics

#### Slice preparation

P70 - 80 mice were deeply anesthetized with isoflurane and quickly decapitated. Acute coronal slices (300 μm) were obtained using a vibrating blade microtome (Leica V1200S) in ice-cold N-Methyl D-Glucamine (NMDG) cutting solution: containing (in mM): 110 NMDG, 20 HEPES, 25 glucose, 30 NaHCO3, 1.25 NaH2PO4, 2.5 KCl, 5 ascorbic acid, 3 Na-pyruvate, 2 Thiourea, 10 MgSO4-7 H_2_O, 0.5 CaCl2, 305-310 mOsm, pH 7.4. Slices equilibrated in a homemade chamber for 2 – 3 min (31°C) in the above solution and an additional 60 min in room temperature aCSF containing (in mM): 120 NaCl, 26 NaHCO3, 1 NaH2PO4, 2.5 KCl, 11 Glucose, 1.3 MgSO4-7 H_2_O, and 2.5 CaCl2 (290-300 mOsm, pH 7.4) before being transferred to a recording chamber.

#### Whole-Cell Patch-Clamp

Recordings were obtained from medium spiny neurons (MSNs) in the dorsal and ventral striatum. Striatal MSNs were visualized under IR-DIC. Data were collected with a Multiclamp 700B amplifier, Digidata 1550B (Molecular devices), and using Clampex 11 (pClamp; Molecular Devices, San Jose, CA). All recordings were acquired in voltage clamp (V_h_ = -70 mV) at 31°C and QX-314 (1uM) was used in all internal solutions to internally block sodium-channels. Whole cell currents were acquired and sampled at 10 kHz with a lowpass bessel filter set at 2 kHz and digitized at 10 kHz. For excitatory currents (EPSCs), the patch pipette was filled with internal solution containing (in mM): 135 CsMeSO4, 8 CsCl, 10 HEPES, 0.25 EGTA, 10 Phosphocreatine, 4 MgATP, and 0.3 NaGTP (295 – 305 mOsm, pH 7.2 with CsOH) and Picrotoxin (50μM) was added to aCSF. For inhibitory postsynaptic currents (IPSC), patch pipettes were filled with internal solution containing (in mM): 143 CsCl, 10 HEPES, 0.25 EGTA, 10 Phosphocreatine, 4 MgATP, and 0.3 Na-2GTP (osmolarity 295 – 305, pH 7.2 with CsOH) and CNQX (50μM) and AP5 (10μM) were added to aCSF. All pipettes (3 - 4 MΩ) were pulled from borosilicate glass (Narishige PC-100). Patched cells were allowed a minimum of 3 min to stabilize following break-in and access resistance (Ra) was monitored throughout the recording and cells with an increase of > 20% in Ra were discarded. Optically evoked synaptic currents were induced with 440nm wavelength LED light delivered through a 40X objective lens (Olympus BX51WI) at 0.1 Hz (5 ms pulse) and light intensity was adjusted using a LLE-SOLA-SE2 controller (Lumencore). Electrically evoked synaptic currents in MSNs were evoked using a bipolar stimulation electrode placed in the dorsal or ventral striatum using the following parameters: 0.1 Hz frequency, pulse duration 0.1ms. Input/output (I/O) curves were also obtained to compare synaptic strength. They were measured with a stimulation at a frequency of 0.1Hz with intensity increments of 10µA (starting at 10µA and finishing at 100µA) with 3 sweeps per increment. We also measured paired-pulse ratio (PPR) to examine presynaptic probability of neurotransmitter release. PPR of evoked EPSCs and IPSCs were obtained with 40, 50 and 100ms inter-pulse intervals and by dividing the amplitude of the second EPSC or IPSC by the first one (EPSC2/EPSC1 or IPSC2/IPSC1). Spontaneous postsynaptic currents (sEPSCs and sIPSCs) were analyzed from epochs between sweeps of optically evoked currents. Events were detected automatically using Clampfit 11 and amplitude threshold for event detection were set at -10 pA and -15 pA for sEPSC and sIPSC, respectively.

#### Immunohistochemistry on brain slices

Nrxn123 WT and Nrxn123 KO mice (P90) were anesthetized using pentobarbital NaCl solution (7 mg/mL) injected intraperitoneally and then perfused intracardially with 20 mL of PBS followed by 30 mL of 4% PFA. The brains were extracted, placed in PFA for 48h and then in a 30% sucrose solution for 48h. After this period, brains were frozen in -30°C isopentane for 1 min. 40 microns thick coronal sections were then cut using a cryostat (Leica CM1800) and placed in antifreeze solution at -20°C until used. For slice immunostaining, after a PBS wash, the tissue was permeabilized, nonspecific binding sites were blocked and slices were incubated overnight with a rabbit anti-TH (1:1000, AB152, Millipore Sigma, USA), a mouse anti-TH (1:1000, Clone LNC1, MAB318, Millipore Sigma, USA), a rat anti-DAT (1:2000, MAB369; MilliporeSigma, USA), a rabbit anti-VMAT2 (1:2000, kindly provided by Dr. G.W. Miller, Columbia University) or a chicken anti-GFP (1:1000, GFP-1020; Aves Labs, USA) antibody. Primary antibodies were subsequently detected with rabbit, rat or chicken Alexa Fluor-488–conjugated, 546-conjugated and/or 647-conjugated secondary antibodies (1:500, 2h incubation; Invitrogen). Slices were mounted on charged microscope slides (Superfrost/Plus, Fisher Scientific, Canada) and stored at 4°C prior to image acquisition.

#### Fast scan cyclic voltammetry

Nrxn123 WT, Nrxn123 HET and Nrxn123 KO mice (P90) were used for Fast Scan Cyclic Voltammetry (FSCV) recordings. The animals were anesthetized with halothane, quickly decapitated and the brain harvested. Next, the brain was submersed in ice-cold oxygenated artificial cerebrospinal fluid (aCSF) containing (in mM): NaCl (125), KCl (2.5), KH2PO4 (0.3), NaHCO3 (26), glucose (10), CaCl2 (2.4), MgSO4 (1.3) and coronal striatal/nucleus accumbens brain slices of 300µm were prepared with a Leica Vt_10_00S vibrating blade microtome. Once sliced, the tissue was transferred to oxygenated aCSF at room temperature and allowed to recover for at least 1h. For recordings, slices were put in a custom-made recording chamber superfused with aCSF at 1 ml/min and maintained at 32°C with a TC-324B single channel heater controller. All solutions were adjusted at pH 7.35-7.4, 300 mOsm/kg and saturated with 95% O2-5% CO2 at least 30-min prior experiment. Electrically-evoked action potential-induced DA overflow was measured by FSCV using a 7µm diameter carbon-fiber electrode crafted as previously described (Martel et al., 2011) and placed into the tissue ∼100µm below the surface and a bipolar electrode (Plastics One, Roanoke, VA, USA) was placed ∼200µm away. The electrodes were calibrated with 1µM DA in aCSF before and after each recorded slice and the mean of the current values obtained were used to determine the amount of released DA. After use, electrodes were cleaned with isopropyl alcohol (Bioshop). The potential of the carbon fiber electrode was scanned at a rate of 300 V/s according to a 10ms triangular voltage wave (−400 to 1000 mV vs Ag/AgCl) with a 100ms sampling interval, using a headstage preamp (Axon instrument, CV 203BU) and a Axopatch 200B amplifier (Axon Instruments, Union City, CA). Data were acquired using a digidata 1440a analog to digital board converter (Axon Instruments) connected to a personal computer using Clampex (Axon Instruments). Slices were left to stabilize for 20-min before any electrochemical recordings. Evaluation of DA release was achieved by sampling 4 different subregions of the dorsal striatum and 4 different subregions of the ventral striatum (nucleus accumbens core and shell) using slices originating from +1.34 to +0.98 using bregma as a reference. After positioning of the bipolar stimulation and carbon fiber electrodes in the tissue, single pulses (400µA, 1ms) were applied to trigger DA release. After sampling of DA release, per pulse ratio experiments were conducted using one spot of the dorsal striatum and one in the nucleus accumbens core. In each spot, a series of single pulses every 2-min for 10-min was collected as a baseline, follow by a triplicate of single-pulse stimuli intercalated with paired pulses (100 Hz) every 2-min (double pulse of 1ms, 400µA, with an inter-pulse interval of 100ms). DA release was analyzed as the peak height of DA concentrations (rise time) and DA reuptake was determined from the clearance rate of DA which was assumed to follow Michaelis-Menten kinetics. The time constant Tau (t) was used as an uptake parameter and was calculated based on an exponential fitting applied to all traces with a homemade MATLAB script. The paired pulse ratio was calculated using the peak height of the 1^st^ and 2^nd^ pulse as: (2^nd^ pulse - 1^st^) /1^st^.

#### Behavioral testing

Before behavioral experiments, mice were transferred from the colony and were housed with a maximum of 4 mice per cage. All mice were handled for 3 consecutive days prior to start of the different tests. All tests were performed in the same order as described below. The animals were tested between 10:00AM and 4:30PM.

#### Rotarod

The accelerating rotarod task was used to assess motor coordination and learning. The apparatus consisted of five rotating rods separated by walls and elevated 30 cm from the ground. P60-P70 mice were pre-trained on the rod (LE8205, Harvard apparatus) one day before the recording to reach a stable performance. Mice were required to remain on the rod for 1-min at a constant speed of 4rpm with a maximum of 3 attempts. For the first step of the rotarod testing protocol, the first day of the data acquisition, mice were tested on an accelerated rotation 4-40rpm over a 10-min period for two sessions with an interval of 1h. The latency to fall was recorded. The same parameters were used on the second test day, but 3 sessions were performed. On the last day of data acquisition, the mice performed 4 sessions with the same previous parameters. A second protocol was also used, in which mice were tested with an accelerated rotation 4-40rpm over a 2-min period for all sessions with an interval of 1h. Each trial per day was analyzed separately and compared between the genotypes.

#### Locomotor activity and psychostimulant-induced locomotor activity

To evaluate motor behavior, mice were placed in cages (Omnitech Electronics, Inc; USA) designed for activity monitoring using an infrared actimeter (Superflex sensor version 4.6, Omnitech Electronics; 40 x 40 x 30cm) for 20-min. Next, 0.9% saline or the drug treatments were injected intraperitoneally (10ml/kg) in a randomized order for the different genotypes. Horizontal activity was scored for 40-min following the injection. To evaluate psychostimulant-induced motor behaviors, the mice were placed in the infrared actimeter cages (Superflex sensor version 4.6, Omnitech Electronics) for 20-min. Then, amphetamine was injected intraperitoneally at 5-mg/kg (Tocris, UK) or cocaine hydrochloride at 20-mg/Kg (Medisca, cat# 53-21-4, Canada) in a randomized order for the different genotypes. The total distance (horizontal locomotor activity) was scored for 40-min following the injection.

#### Pole test

The test was conducted with a homemade 48-cm metal rod of 1-cm diameter covered with adhesive tape to facilitate traction, placed in a cage. 8-week-old Nrxn123 KO, HET and WT mice were positioned head-up at the top of the pole and the time required to turn (t-turn) and climb down completely was recorded.

#### Sucrose preference test

Mice were tested for preference of a 2% sucrose solution, using a two bottles choice procedure. Subjects were housed one per cage all of the test (5 days). Mice were first conditioned with two bottles of water during two days. Then mice were given two bottles, one sucrose (2%) and one of tap water. Every 24h, the amount of sucrose and water consumed was evaluated. To prevent potential location preference of drinking, the position of the bottles was changed every 24h. The preference for the sucrose solution was calculated as the percentage of sucrose solution ingested relative to the total intake.

### Reverse transcription quantitative polymerase chain reaction (RT-qPCR)

We used RT-qPCR to quantify the amount of mRNA encoding the following genes: gephyrin (Gphn), collybistin (Arhgef 9), GABA-A receptor (Gabra1), neuroligins 1, 2 and 3 (Nlgn1, 2 and3), latrophilins 1, 2 and 3 (Lphn1, 2 and 3), LRRTMs 1, 2, 3, 4 (LRRTM1, 2, 3 and 4), D1R (DRD1) and D2R (DRD2) in striatal brain tissue from P80 Nrxn123 WT and Nrxn123 KO mice. The brains were quickly harvested and the ventral and dorsal striata were microdissected and homogenized in 500 µL of trizol. For both, presynaptic and postsynaptic compartment, RNA extraction was performed using RNAeasy Mini Kit (Qiagen, Canada) according to the manufacturer’s instructions. RNA integrity was validated using a Bioanalyzer 2100 (Agilent). Total RNA was treated with DNase and reverse transcribed using the Maxima First Strand cDNA synthesis kit with ds DNase (Thermo Fisher Scientific). Gene expression was determined using assays designed with the Universal Probe Library from Roche (www.universalprobelibrary.com). For each qPCR assay, a standard curve was performed to ensure that the efficiency of the assay was between 90% and 110%. The primers used are listed in **supplementary Table 1**. The Quant Studio qPCR instrument (Thermo Fisher Scientific) was used to detect the amplification level. Relative expression comparison (RQ = 2~^-ΔΔCT^) was calculated using the Expression Suite software (Thermo Fisher Scientific), using GAPDH and β-Actin as an endogenous control.

### Western Blot

Ventral and dorsal striatum samples were microdissected from Nrxn123 WT and Nrxn123 KO adult mice and then lysed in RIPA buffer (Fisher Scientific, PI89900) containing a protease inhibitor cocktail (P840, Sigma). Homogenized tissue samples were centrifuged at 12000g for 30 min at 40C. Supernatant was collected, and protein quantification was done with BCA reagent (Thermo Scientific Pierce BCA Protein Assay Kit, PI23227). 20μg of each sample was separated on 8% SDS-PAGE followed by transfer onto a nitrocellulose membrane. Membrane blocking was done with 10% skimmed milk for 90 min at RT with gentle shaking. The membranes were incubated overnight at 4°C with respective antibodies (**Supplementary Table 2**) with gentle shaking. Membranes were washed 5 times with TBST buffer for 5 min each time. After this, appropriate secondary antibodies (**Supplementary Table 2**) were added, and the incubation was performed at RT for 90 min with gentle shaking. Membranes were washed again with TBST buffer for 5 times X 5 min and developed using the Clarity Western ECL substrate (Bio-Rad, 1004384863). Images were captured on a luminescent image analyzer (GE Healthcare) using Image quant LAS 4000 software. Membranes were stripped and re-probed for β-actin as a loading control.

### Image acquisition with confocal microscopy

All *in vitro* fluorescence imaging quantification analyses were performed on images acquired using an Olympus Fluoview FV1000 point-scanning confocal microscope (Olympus, Tokyo, Japan).

Images were scanned sequentially to prevent non-specific bleed-through signal using 488, 546 and 647-nm laser excitation and a 60X (NA 1:42) oil immersion objective.

### Image analysis

All image quantification was performed using Image-J (National Institute of Health, NIH) software. A background correction was first applied at the same level for every image analyzed before any quantification. A macro developed in-house was used to perform all quantifications.

### Statistics

Data are represented throughout as mean ± SEM. Statistically significant differences were analyzed using Student’s t test, one-way repeated measures ANOVA or two-way ANOVA with Tukey’s or Sidak’s multiple comparison test (*p < 0.05; **p < 0.01; ***p < 0.001; ****p < 0.0001).

## Acknowledgments

We would like to thank Dr. G. Miller for kindly provided VMAT2 antibody, Willemieke Kouwenhoven for her help with some experiments, the IRIC genomic platform for qRT-PCR analysis, CERVO Québec electron microscopy platform and C.A. Maurice for his strong support. This work was funded by Canadian Institutes of Health Research (CIHR, grant MOP106556) to L-E.T. C.D. received a graduate student award from Fond de recherche en santé du Québec (FRQS).

## Author contributions

C.D.: conceptualization, design, acquisition, analysis, validation and interpretation of data (All *in vitro* results, ICC, GABA uptake, Behavior, IHC, qRT-PCR, Transmission electron microscopy), drafting, revising and editing the manuscript. G. D. C: design, acquisition of data, analysis (whole cell patch clamp recording, drafting, revising and editing the manuscript), validation, and interpretation of data. C. VL. D.: acquisition of TEM. S.M.: acquisition and analysis of data (Western Blot). N.G.: viral injections. S.B.N.: help with data analysis (ICC). M-J.: acquisition of data (cell culture), B. D. L.: acquisition of data and analysis (voltammetry). L. Y. C.: conceptualization, design, resources, providing Nrxn123flox/flox mice, genotyping and injection verification, electrophysiology analysis, writing, and review and editing. L-E.T.: conceptualization, resources, funding acquisition, writing, review and editing.

## Supplementary figures and legends

**Supplementary figure 1.**
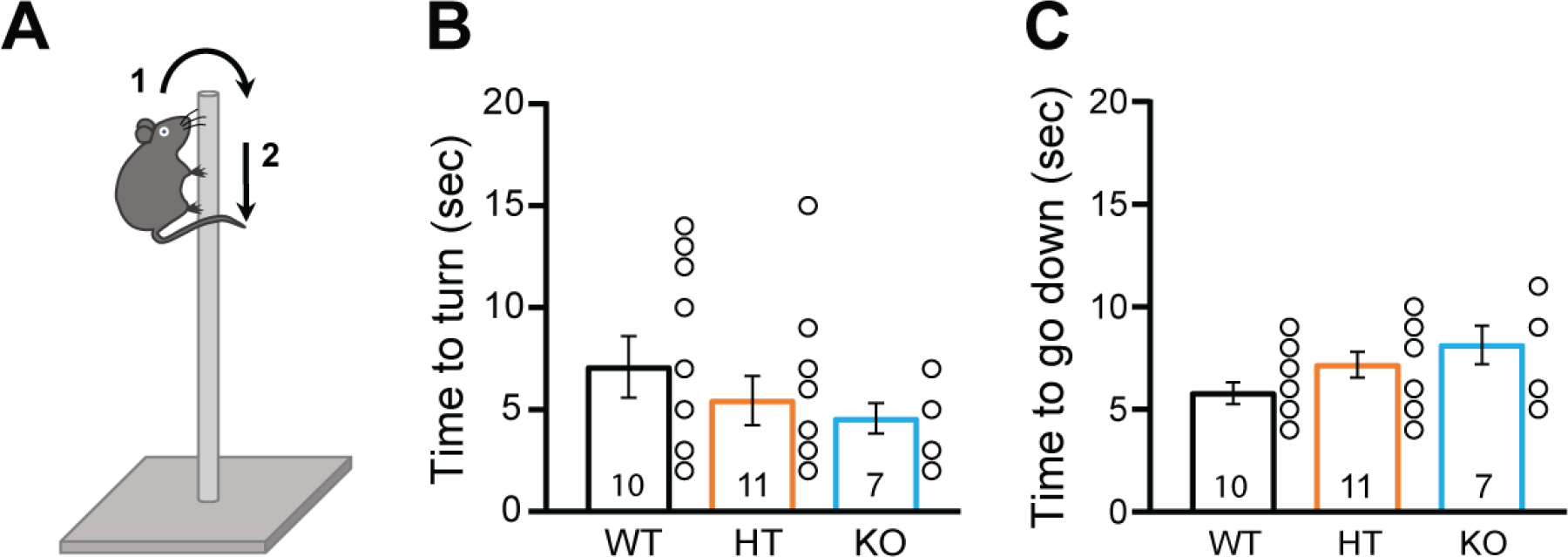
No motor deficit with the pole test in DAT::NrxnsKO mice. **A**- schematic representation of the pole test procedure. **B**- Summary graph of the time to turn shows no genotype effect (WT= 7.10 ± 1.50sec; HET= 5.45 ± 1.20sec; KO= 4.57 ± 0.75sec). **C**- Summary graph of the time to go down shows no genotype effect (WT= 5.80 ± 0.53sec; HET= 7.18 ± 0.62sec; KO= 8.14 ± 0.93sec) (WT n=10; HET n=11 and KO n=7 mice).

**Supplementary figure 2.**
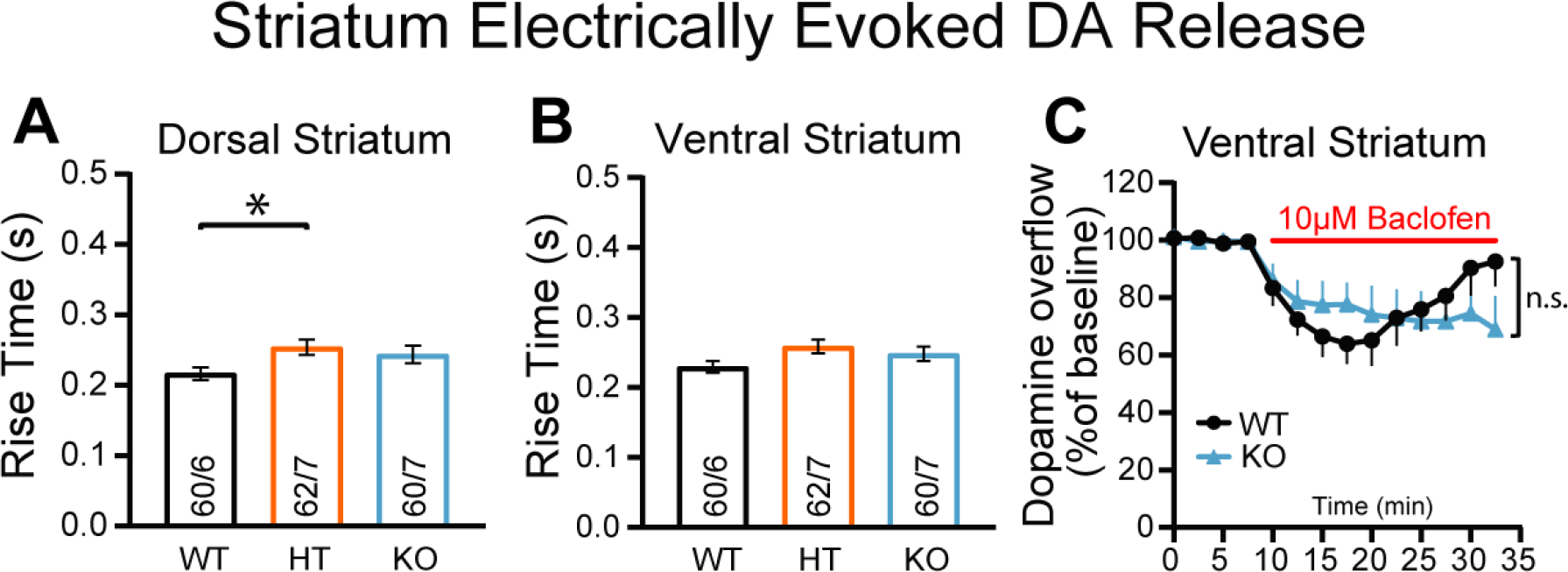
No change in GABA_B_ receptor modulation of DA release after conditional deletion of all neurexins. **A-** Summary graph of the rise time of electrically evoked DA overflow in the dorsal striatum showing a genotype difference (HET versus WT : 0.25 ± 0.01s compared to 0.22 ± 0.01s, respectively). **B**- Summary graph of the rise time of electrically evoked DA overflow in the ventral striatum shows no genotype effect. **C**- Plot of relative peak DA overflow and its modulation by the GABA_B_ agonist baclofen in the ventral striatum. No difference between WT and KO mice was observed.

**Supplementary figure 3.**
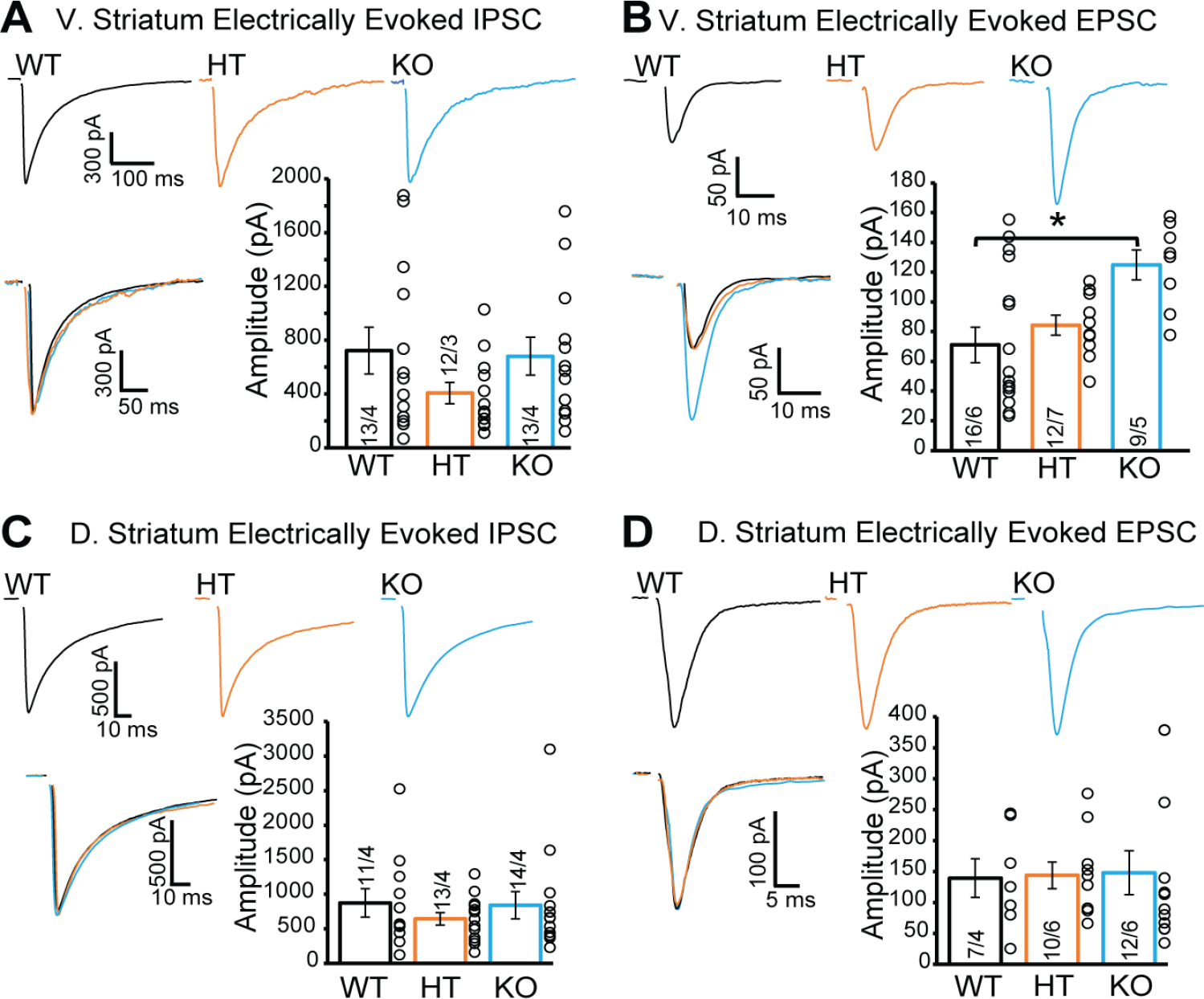
Electrically evoked EPSC amplitude is increased in the vSTR of DAT::NrxnsKO mice. **A-** Representative traces (top) of IPSCs elicited by electrical-stimulation (e-IPSC) and recorded in ventral striatal MSNs. Summary graph of e-IPSC amplitudes (bottom) shows no genotype effect (*Peak Amplitude:* WT = -722.64 ± 174.29 pA, HT = -406.01 ± 80.15 pA, KO = -680.89 ± 141.06, Mean ± SEM). **B-** Representative traces (top) of EPSCs elicited by electrical-stimulation (e-EPSC) and recorded in ventral striatal MSNs. Summary graph of e-EPSC amplitudes (bottom) shows an increased amplitude in the KO mice compared to WT (Peak Amplitude: WT = -71.03 ± 11.9 pA, HT = -84.2 ± 6.76 pA, KO = -124.8 ± 10.15, Mean ± SEM). **C**- Representative traces (top) e-IPSCs recorded in dorsal striatal MSNs. Summary graph of e-IPSC amplitudes (bottom) shows no genotype effect. **D**- Representative traces (top) of e-EPSCs recorded in dorsal striatal MSNs. Summary graph of e-EPSC amplitudes (bottom) shows no genotype effect. Data in summary graphs are presented as mean ± SEM; statistical comparisons were performed with One-Way ANOVA (*P<0.05; **P<0.01; ***P<0.001; non-significant comparisons are not identified). Tukey’s correction was used for all multiple comparisons.

**Supplementary figure 4.**
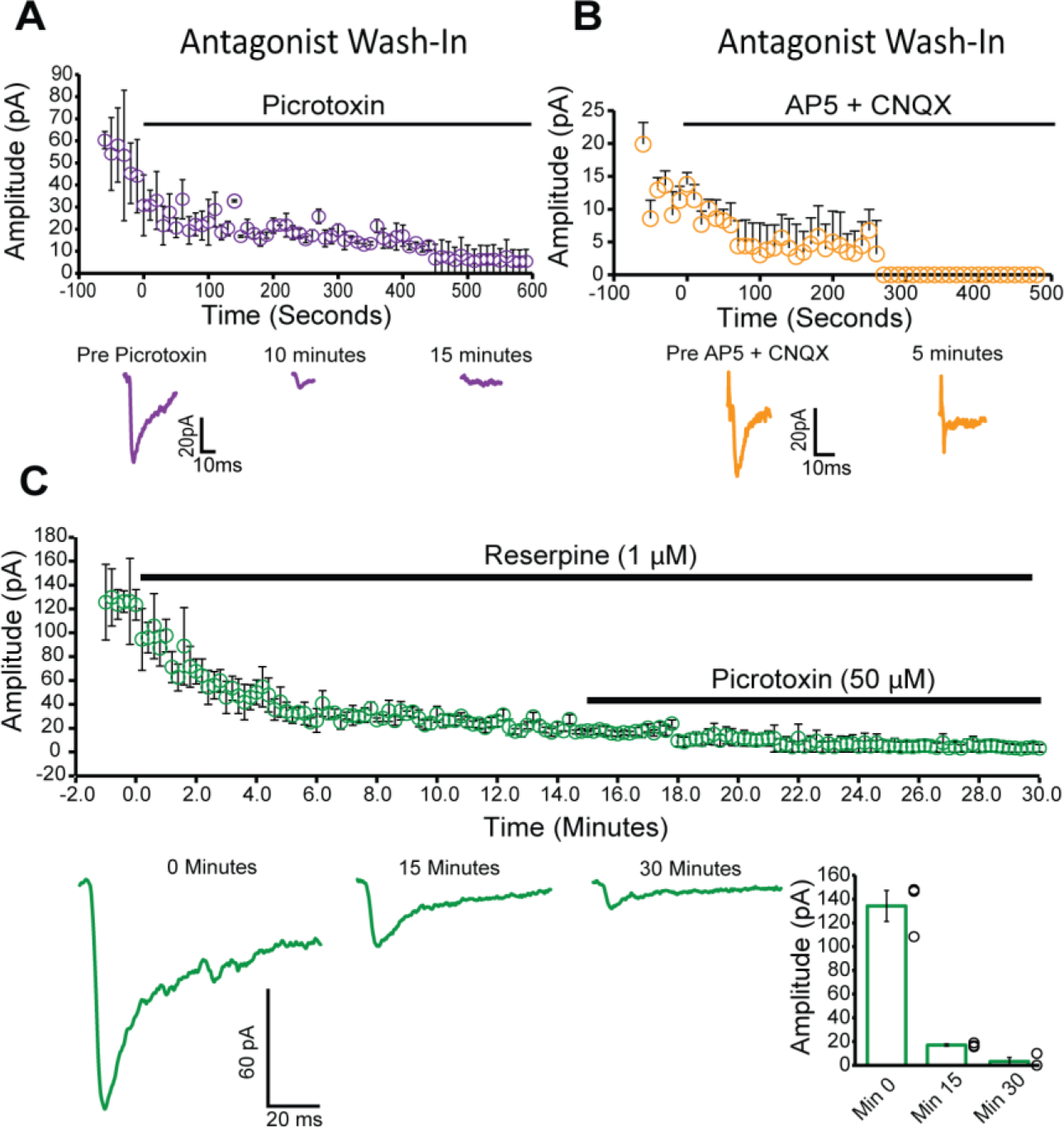
Pharmacological properties of o-IPSCs and o-EPSCs recorded from MSNs in acute ventral striatal slices. **A**- Graph showing the amplitude of o-**IPSCs** recorded in ventral striatal MSNs before and after incubation with the GABA_A_ channel receptor pore blocker Picrotoxin. **B**- Graph showing the amplitude of o-EPSCs recorded in ventral striatal MSNs before and after incubation with the AMPA and NMDA receptor blockers AP5 and CNQX, respectively. **C**-Graph showing the amplitude of o-IPSCs recorded in ventral striatal MSNs before and after incubation with the VMAT2 blocker reserpine, followed by Picrotoxin.

**Supplementary figure 5.**
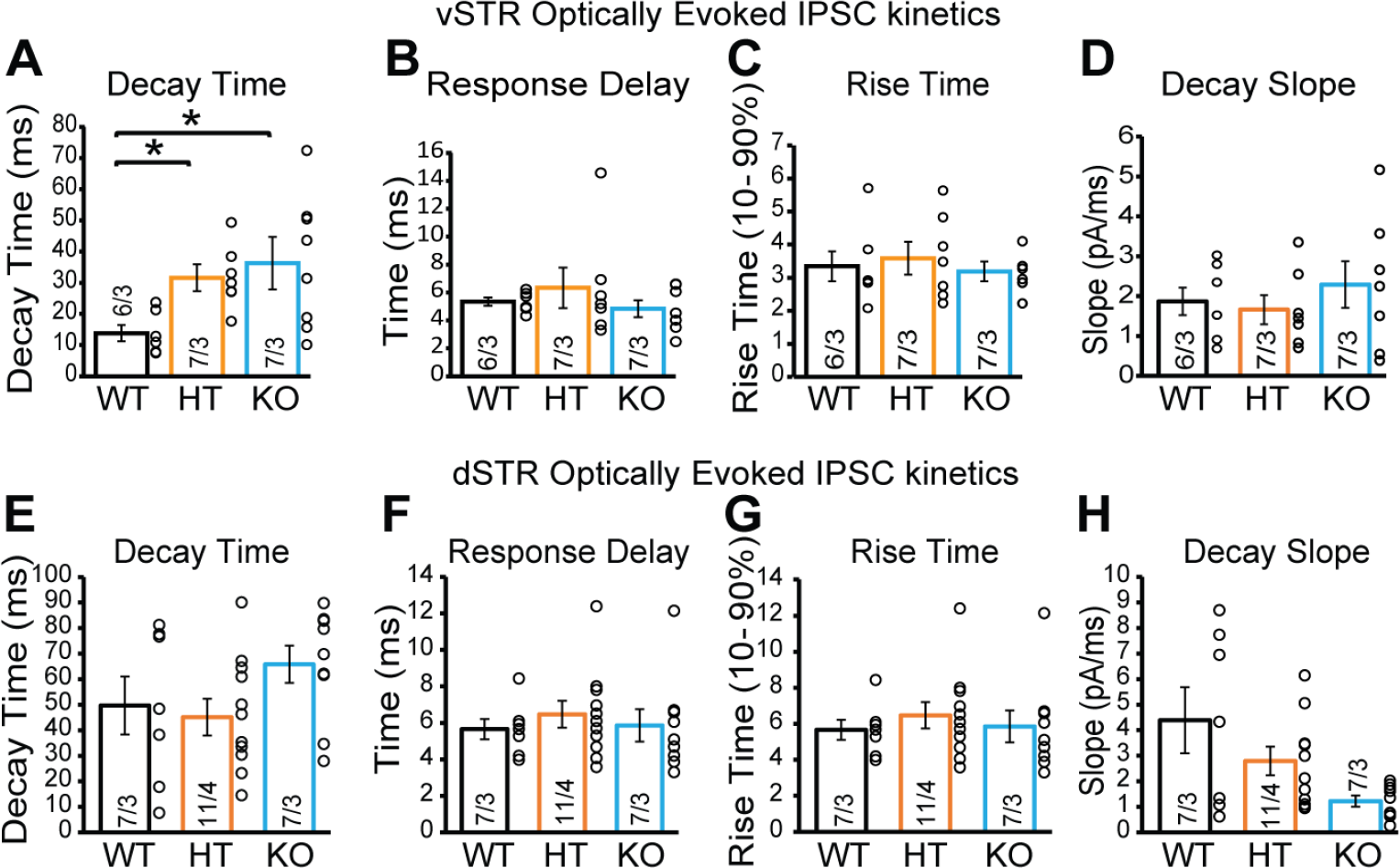
Removal of neurexins increases oIPSC decay time in the ventral but not the dorsal striatum. **A to D -** Summary graphs of o-IPSC kinetics recorded in ventral striatal MSNs. From left to right: decay time (**A**) (Time [90 – 10%]: WT = 13.8 ± 2.5 ms, HT = 31.6 ± 4.3 ms, KO = 36.2 ± 8.4 ms, Mean ± SEM), response delay (latency from stimulation to peak response) (**B**), rise time (**C**), and decay slope (**D**). **E to H -** Summary graphs of o-IPSC kinetics recorded in dorsal striatal MSNs; (from left to right) decay time (**E**), response delay (**F**), rise time (**G**), and decay slope (**H**).

**Supplementary figure 6.**
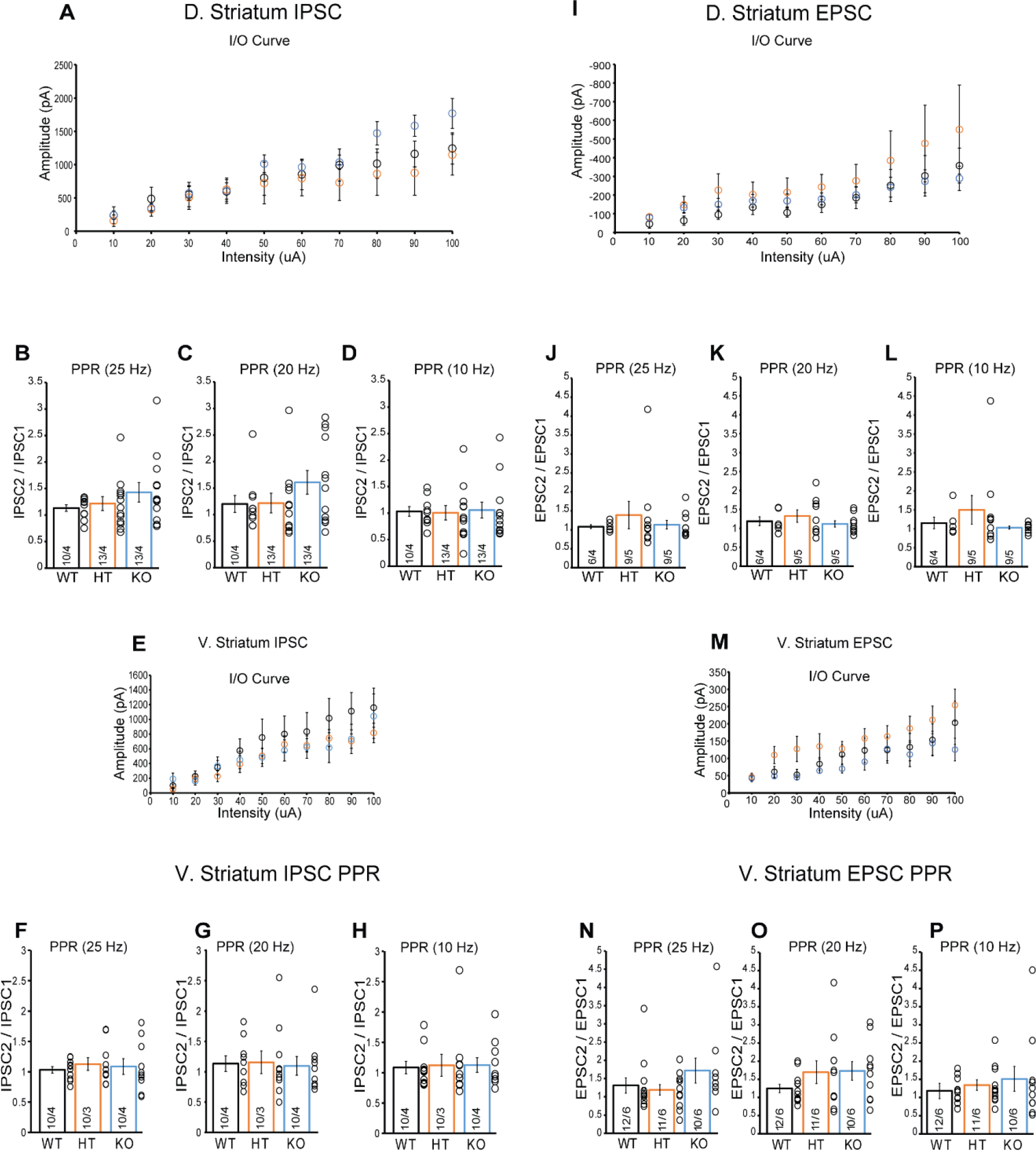
No change in synaptic plasticity of electrically-evoked IPSCs and EPSCs after conditional deletion of all Nrxns in DA neurons. - **A.** Scatterplot of IPSC input- output relationship in MSNs of the dorsal striatum of WT, HT, and KO mice. **B.** Summary graph of GABAergic IPSC paired-pulse ratio (PPR) in MSNs of the dorsal striatum; two electrical pulses were delivered with an inter-stimulation interval (ISI) of 40 ms (25 Hz). **C.** Same as in **B**; ISI = 50 ms (20 Hz). **D.** Same as in **B** and **C**; ISI = 100 ms (10 Hz). **E.** Scatterplot of IPSC input-output relationship in MSNs of the ventral striatum. **F.** Summary graph of GABAergic IPSC paired-pulse ratio (PPR) in MSNs of the ventral striatum; two electrical pulses were delivered with an inter-stimulation interval (ISI) of 40 ms (25 Hz). **G.** Same as in **F**; ISI = 50 ms (20 Hz). **H.** Same as in **F** and **G**; ISI = 100 ms (10 Hz). **I.** Scatterplot of EPSC input-output relationship in MSNs of the dorsal striatum. **J.** Summary graph of glutamatergic paired-pulse ratio (PPR) in MSNs of the dorsal striatum; two electrical pulses were delivered with an inter-stimulation interval (ISI) of 40 ms (25 Hz). **K.** Same as in **J**; ISI = 50 ms (20 Hz). **L.** Same as in **K** and **J**; ISI = 100 ms (10 Hz). **M.** Scatterplot of EPSC input-output relationship in MSNs of the ventral striatum. **N.** Summary graph of EPSC paired-pulse ratio (PPR) in MSNs of the ventral striatum; two electrical pulses were delivered with an inter-stimulation interval (ISI) of 40 ms (25 Hz). **O.** Same as in **N**; ISI = 50 ms (20 Hz). **P.** Same as in **N** and **O**; ISI = 100 ms (10 Hz).

**Supplementary figure 7.**
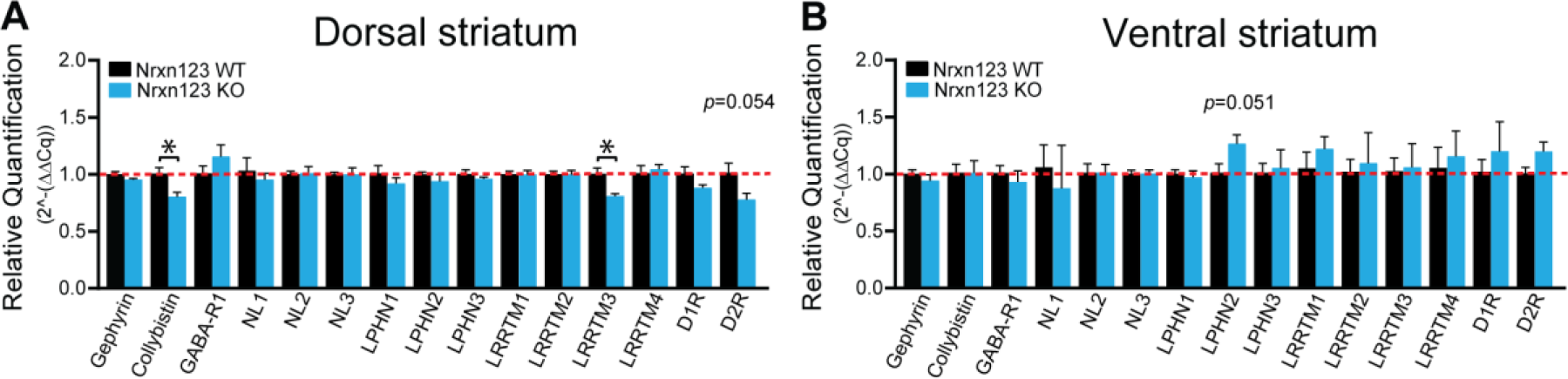
Gene expression profile in target cells of DA neurons after conditional deletion of Nrxn123. **A** and **B**– Relative changes of mRNA levels measured by RT-qPCR, from ventral and dorsal striatum tissue: Gephyrin (Gphn), Collybistin (Arhgef 9), GABA-A receptor (Gabra1), Neuroligins 1, 2 and 3 (Nlgn1, 2 and 3), Latrophilins 1, 2 and 3 (Lphn1, 2 and 3), LRRTMs 1, 2, 3, 4 (LRRTM1, 2, 3 and 4), D1R (DRD1) and D2R (DRD2) in brain tissue from P80 DAT::NrxnsWT, and DAT::NrxnsKO mice.

**Supplementary figure 8.**
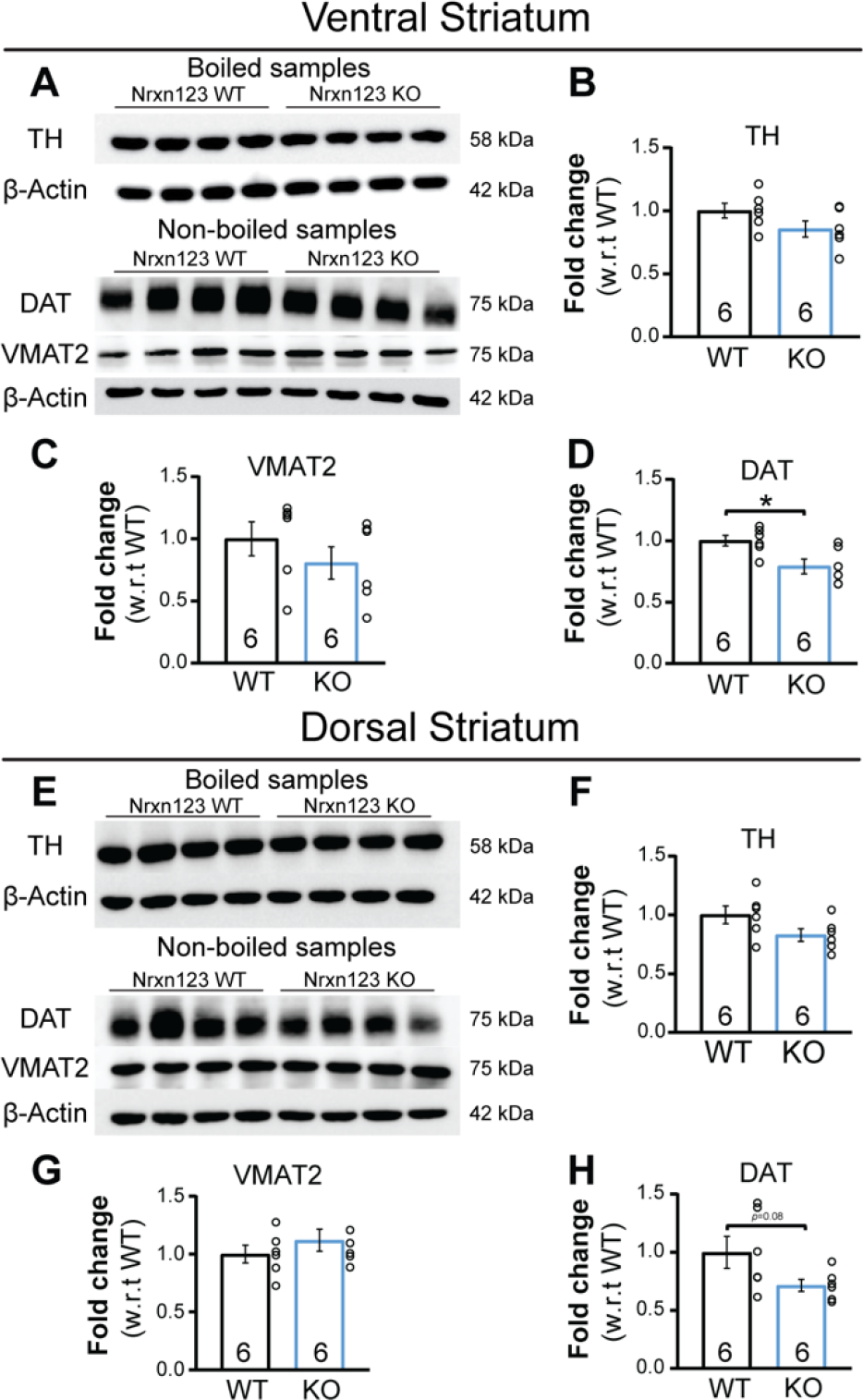
Decrease of DAT protein expression in DA neurons from DAT::NrxnsKO mice. **A-** Representative western blots illustrating TH, DAT, VMAT2 and β-actin protein from total striatum homogenates of adult mice. **B to D**- Immunoblot quantifying relative protein levels (fold change compared to controls) for TH (**B**), VMAT2 (**C**) and DAT (**D**) (n = 6 DAT::NrxnsWT, HET and KO mice). **E-** Representative western blots illustrating TH, DAT, VMAT2 or β-actin from total striatum homogenates of adult mice. **F to H**- Immunoblot quantifying relative protein levels (fold change compared to controls) for TH (**F**), VMAT2 (**G**) and DAT (**H**) (n = 6 DAT::NrxnsWT, HET and KO mice). Error bars represent ± S.E.M. (ns, non-significant; *, p < 0.05; **, p < 0.01; ***, p < 0.001; ****, p < 0.0001).

**Supplementary table 1.**
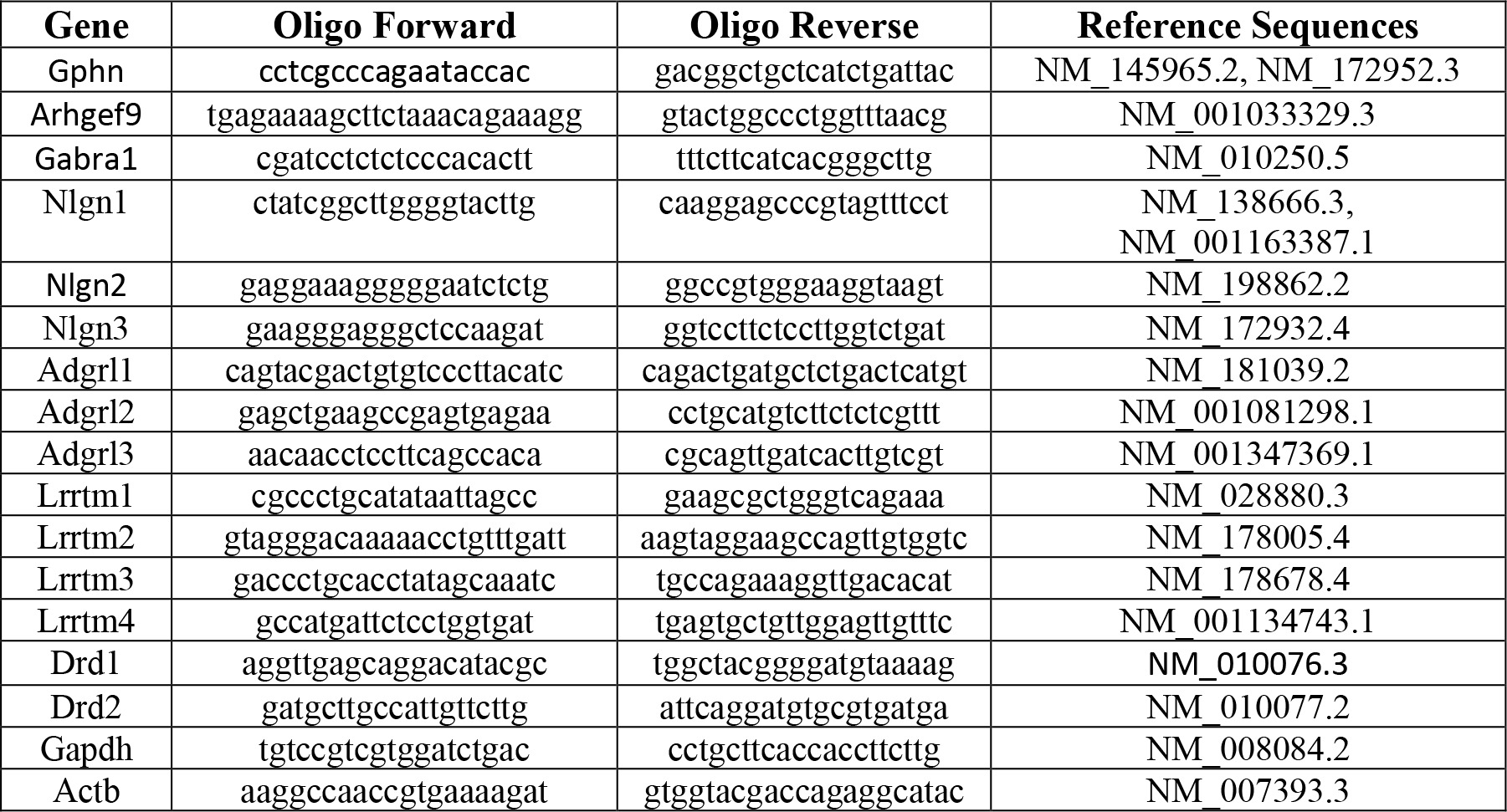
qPCR primers for ventral and dorsal striatum.

**Supplementary table 2.** Antibodies for Western Blot.

| **Antibody** | **Catalog numbers** | **Raised in** | **Dilution used** | **RRID** |
| --- | --- | --- | --- | --- |
| DAT | MAB369<br>(Millipore) | Rat | 1:1000 | AB_2190413 |
| TH | MAB318<br>(Millipore) | Mouse | 1:1000 | AB_2201528 |
| VMAT2 | Kind gift from Dr. Gary Miller,<br>Emory University, USA | Rabbit | 1:5000 | NA |
| β-Actin | A3854(Sigma) | Purified from<br>hybridoma<br>cell culture | 1:15000 | AB_262011 |

| **Antibody** | **Catalog number** | **Dilution used** | **RRID** |
| --- | --- | --- | --- |
| Peroxidase-conjugated<br>AffiniPure Goat Anti-Rat<br>IgG (H+L) | 711-035-152<br>(Cedarlane) | 1:5000 | AB_10015282 |
| Peroxidase-conjugated<br>AffiniPure Goat Anti-Rabbit<br>IgG (H+L) | 112-035-003<br>(Cedarlane) | 1:5000 | AB_2338128 |
| Peroxidase-conjugated<br>AffiniPure Goat Anti-Mouse<br>IgG (H+L) | 115-035-146<br>(Cedarlane) | 1:5000 | AB_2307392 |

